# Identifying Context-Specific Cell-Cell Interaction Genes Without Ligand-Receptor Databases from Spatial Transcriptomics

**DOI:** 10.64898/2026.05.08.723913

**Authors:** Hyobin Kim, Beomsu Park, Junghyun Jung, Seulgi Lee, Sanaz Panahandeh, Soyoung Kwon, Jingyi Jessica Li, Esha Madan, Dongha Kim, Junil Kim, Rajan Gogna, Kyoung Jae Won

## Abstract

Current approaches to inferring cell-cell interactions (CCIs) are largely constrained by predefined ligand-receptor databases, particularly for low-resolution spatial transcriptomics (ST) platforms such as Visium. Due to the difficulties in accurately resolving interacting cells at coarse spatial resolution, other modes of interaction are often overlooked. Low-resolution ST data, however, can serve as an alternative to high-resolution ST, which suffers from low sensitivity, and to image-based ST, which is limited by restricted gene panels.

Here, we present CellNeighborEX v2, a database-free framework that directly infers CCI-associated genes from ST data by detecting deviations between observed and expected gene expression at the spot-population level. These deviations are rigorously evaluated through a hybrid statistical framework involving permutation testing and are further refined by considering the abundance of interacting cell-type pairs. Compared with other conventional approaches relying on ligand-receptor databases, CellNeighborEX v2 can capture CCI genes from a broad spectrum of interactions, including both paracrine signaling and contact-dependent communication. Across datasets from hippocampus, liver cancer, colorectal cancer, ovarian cancer, and lymph node infection, CellNeighborEX v2 accurately recapitulated previously identified CCIs. Notably, it uniquely detected interactions absent from existing ligand-receptor databases, enabling detection of context-specific CCIs from Visium data. CellNeighborEX v2 is a tool that expands the analytical spectrum of Visium data and deepens our understanding of the molecular language of intercellular communication.

## Introduction

Understanding how cells communicate within complex tissue environments is essential for elucidating the mechanisms that govern tissue development, immune responses, and disease progression^1-3^. Cell-cell interactions (CCIs) influence these processes through a variety of mechanisms, including paracrine signaling via ligands and receptors, direct cell contact, and mechanical forces transmitted through the extracellular matrix (ECM)^4-6^. Currently, the majority of approaches rely primarily on ligand-receptor expression to detect CCIs, which limits the understanding of other modes of cellular communication. Consequently, the specific genes affected by CCIs and their functional roles remain largely unknown.

The advent of spatial transcriptomics (ST) technologies has enabled transcriptome-wide gene expression measurements in intact tissues, allowing researchers to investigate how local CCIs contribute to gene regulation in situ^7-9^. Yet, significant challenges remain. Image-based platforms like Xenium^10^ offer true single-cell resolution and spatial precision. However, their reliance on targeted gene panels constrains their ability to capture full transcriptome, limiting their utility for discovering unknown CCI genes^11^. In contrast, next generation sequencing (NGS)-based high-resolution technologies such as Visium HD^12^, Slide-seq^13-15^, and Stereo-seq^16,17^ provide significantly higher spatial resolution, enabling finer cellular and subcellular mapping. However, these platforms often suffer from reduced mRNA capture efficiency, particularly for low-abundance transcripts, and are accompanied by increased technical complexity and cost^11,13^. These limitations hinder their widespread adoption, especially in clinical or large-cohort studies. Moreover, due to computational and analytical challenges, data from these technologies are often aggregated into lower-resolution representations for interpretation and further analysis^16-20^.

10X Genomics Visium^7,10^ offers comprehensive coverage and compatibility with diverse sample types including formalin-fixed paraffin-embedded (FFPE) tissues. Its robustness, scalability, and cost-effectiveness make it particularly well-suited for large-scale and diverse biological studies. However, the accessibility of Visium comes at the cost of spatial resolution. Each Visium capture spot spans 55 *µ*m in diameter and collects transcripts from multiple adjacent cells. This inherent process of multicellular averaging blurs cellular boundaries, masking fine-scale spatial gene expression and complicating the detection of CCIs.

Due to the limitations in Visium data, studying CCIs has heavily relied on predefined ligand-receptor databases^21-24^, which fundamentally limits their capacity to discover novel interaction-driven genes beyond known signaling pathways. Interactions between Visium spots have been studied by annotating each spot with its majority cell type^21,23-26^. However, these approaches overlook cellular interactions occurring within individual spots, where multiple cells are in direct proximity or physical contact. While recent studies using physically interacting cells and high-resolution spatial transcriptomics data have identified such interactions^27,28^, no current approach has yet enabled their detection specifically from Visium data. Moreover, existing approaches generally lack the capability to systematically compare interaction patterns across different biological contexts such as spatial regions, tissue subtypes, or experimental conditions^21-26,29-32^.

To address these challenges, we developed CellNeighborEX v2, a database-free computational framework designed to identify a broad spectrum of cell-cell interaction-driven genes (CCI genes) in low-resolution ST data. In this study, CCI genes are defined as transcripts whose expression is significantly associated with the local proximity of two or more cell types within individual spots. Because it does not rely on a ligand-receptor database, it has the potential to detect a wide range of communication modalities, including canonical ligand-receptor signaling, direct cell-cell contact, and extracellular matrix (ECM)-mediated interactions.

CellNeighborEX v2 identifies candidate CCI genes by quantifying deviations between expected and observed gene expression within each Visium spot. For Visium data, expected expression can be modeled using cell type deconvolution results coupled with matched scRNA-seq reference profiles. To obtain CCI gene candidates, CellNeighborEX v2 employs a population-based refinement strategy to Visium spots. After clustering Visium spots according to a biological context, it selects genes exhibiting significantly elevated expression through rigorous statistical testing. To further refine CCI genes, CellNeighborEX v2 uses a regression framework to evaluate whether gene expression is associated with the abundance of candidate interacting cell type pairs. Consequently, CellNeighborEX v2 can capture diverse forms of CCIs.

We conducted a comprehensive evaluation of CellNeighborEX v2 using both synthetic datasets and a wide range of biological samples encompassing various tissue types and disease contexts. In controlled simulations with ground-truth annotations, CellNeighborEX v2 consistently outperformed existing computational methods in recovering genes driven by CCIs. It demonstrated strong performance in detecting spatial region-specific interaction genes that are elevated within defined spatial regions.

To assess its real-world applicability, we applied CellNeighborEX v2 to an extensive panel of biological systems, including mouse hippocampal tissue, mouse hepatocellular carcinoma, human ovarian carcinoma, human colorectal carcinoma, and mouse lymphoid tissue during infection. Across these diverse contexts, the framework reliably identified CCIs that varied across spatial regions, tissue subtypes, and disease states. Notably, the database-free approach of CellNeighborEX v2 enabled the discovery of novel CCI genes from Visium spots that remained undetected by conventional, database-dependent methods. The biological relevance of the inferred interactions was validated through systematic comparison with matched high-resolution spatial transcriptomics datasets, confirming both the accuracy and functional significance of the predictions. In sum, CellNeighborEX v2 is a transformative analytical framework for low-resolution ST data that expands the analytical spectrum to uncover complex CCI mechanisms that were previously inaccessible using conventional computational approaches.

## Results

### Overview of CellNeighborEX v2

CCIs can induce specific gene expression responses in target cells, creating spatially localized and context-dependent communication events. In a tissue, the expression of interaction-associated genes often exhibits selective spatial patterns, with locally enhanced expression levels. This localized upregulation is typically underrepresented in population-level scRNA-seq profiles^27,28^. Similarly, in low-resolution Visium data, these patterns are often obscured as they are averaged across multiple cells within a single capture spot.

CellNeighborEX v2 exploits the complementary strengths of ST and scRNA-seq to detect interaction-driven gene expression. The framework estimates baseline expression expectations for each ST spot by combining scRNA-seq reference profiles with spot-level cell type abundances obtained through deconvolution. While scRNA-seq profiles represent average transcriptional activity across cell populations, their linear combination with deconvolved cell compositions provides expected expression profiles that inherently lack localized, interaction-driven signals. This occurs because CCIs can affect only a fraction of cells within each cell type, and when expression is averaged across all cells of that type in the scRNA-seq reference, these localized signatures become diluted or masked.

CellNeighborEX v2 leverages this fundamental discrepancy to identify CCI genes by systematically comparing observed ST expression levels with scRNA-seq-derived expectations (Fig. 1a). Genes exhibiting consistent positive deviations across relevant biological contexts are classified as putative interaction genes, representing transcripts whose expression is enhanced by cell-cell communication within the intact tissue microenvironment. To illustrate this, we examined *Fabp7*, a CCI gene previously validated via co-culture assays to be specifically induced in astrocytes upon interaction with endothelial tip cells^27^. While its observed ST expression revealed localized enrichment in areas where these cell types co-occur, its expected expression in the scRNA-seq reference remained low due to signal dilution across the entire cell population (Supplementary Fig. 1).

**Fig. 1:**
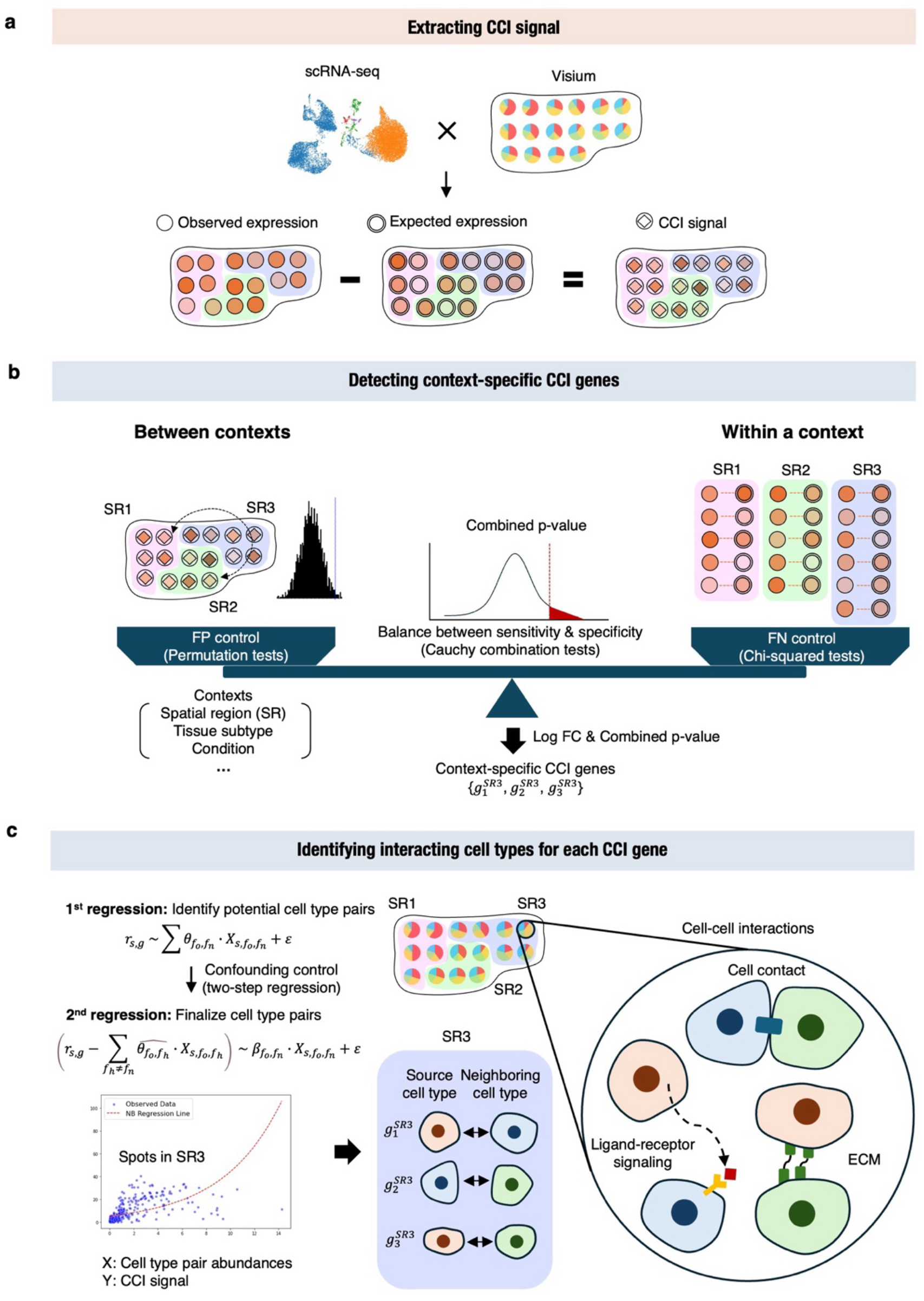
Overview of the CellNeighborEX v2 framework for detecting context-specific cell-cell interaction (CCI) genes. **a** Schematic overview of CellNeighborEX v2. Observed gene expression in Visium data is compared against expected expression levels derived from reference scRNA-seq profiles and deconvolved cell type compositions. Localized overexpression, defined as the difference between observed and expected values, represents CCI signals. **b** Statistical framework for identifying context-specific CCI genes. To capture expression driven by cell-cell interactions, CellNeighborEX v2 integrates a permutation test across contexts and a chi-squired test within a context. P-values from both tests are combined using a Cauchy combination test to balance specificity and sensitivity. **c** Regression-based inference of interacting cell type pairs for each CCI gene. In the first step, a ridge-regularized non-negative negative binomial regression estimates the contribution of candidate source-neighbor cell type pairs to the observed CCI signal across spots. In the second step, a linear model isolates the individual contribution of each pair while controlling for confounding effects. This two-step modeling approach enables directional inference of interaction-driving cell types and captures a range of CCI mechanisms, including ligand–receptor interactions, direct cell–cell contact, and ECM-mediated communication.

To identify CCI genes, CellNeighborEX v2 applies a hybrid statistical testing strategy to clusters defined by biological contexts, accounting for both global and local variation (Methods). Permutation tests across biological contexts such as spatial regions, disease subtypes, or experimental conditions evaluate whether the expression deviations of a gene differ significantly between contexts, effectively controlling for false positives (FPs). In parallel, chi-squared tests within each context assess spatial heterogeneity at the spot level to capture localized expression patterns that may be missed by global comparisons and helps reduce false negatives (FNs). The resulting p-values from the two tests are integrated using a Cauchy combination test^33,34^, balancing sensitivity and specificity to identify a robust and interpretable set of context-specific CCI genes (Fig. 1b). To enable a fair combination of the two tests, permutation-derived p-values were converted from empirical null distributions to Z-scores and then transformed into continuous p-values (Methods). This procedure resolves the discreteness of permutation-derived p-values, allowing direct comparison with the continuous chi-squared p-values.

To further filter CCI genes, CellNeighborEX v2 utilizes cell type abundance of the interacting pairs. CellNeighborEX v2 infers the specific interacting cell type pairs that contribute to the expression of each CCI gene using a two-step regression framework (Methods). In the first step, a ridge-regularized non-negative negative binomial model estimates the contributions of selected candidate cell type pairs to the expression of a given CCI gene. In the second step, a simplified linear model isolates the effect of each individual pair by adjusting for confounding interactions. Each interacting cell type pair is modeled as a directional relationship, where the source cell expresses the CCI gene and the neighboring cell modulates its expression through local interactions (Fig. 1c). This allows CellNeighborEX v2 to function independently of predefined ligand-receptor databases and to detect CCI genes arising from diverse communication modalities including novel signaling pathways, metabolic exchanges, mechanical interactions, ECM and contact-dependent mechanisms that would otherwise remain undetected by conventional database-dependent approaches.

### CellNeighborEX v2 accurately recovers context-specific CCI genes in a synthetic benchmark dataset

To evaluate the performance of CellNeighborEX v2 in detecting CCI genes, we generated a synthetic Visium ST dataset designed to mimic realistic tissue architecture and cell composition (Methods). Spot locations and the total cell numbers per spot were derived from human ovarian cancer Visium data^35^. To estimate the number of cells within each spot, we used annotated scRNA-seq data^35^ from the matching source to perform spot deconvolution (Fig. 2a). For each synthetic spot, total cell counts were sampled from the 5th to 95th percentile of the distribution observed in real Visium data, producing a cell abundance profile that closely matched the original Visium dataset (Methods) (Fig. 2b).

**Fig. 2:**
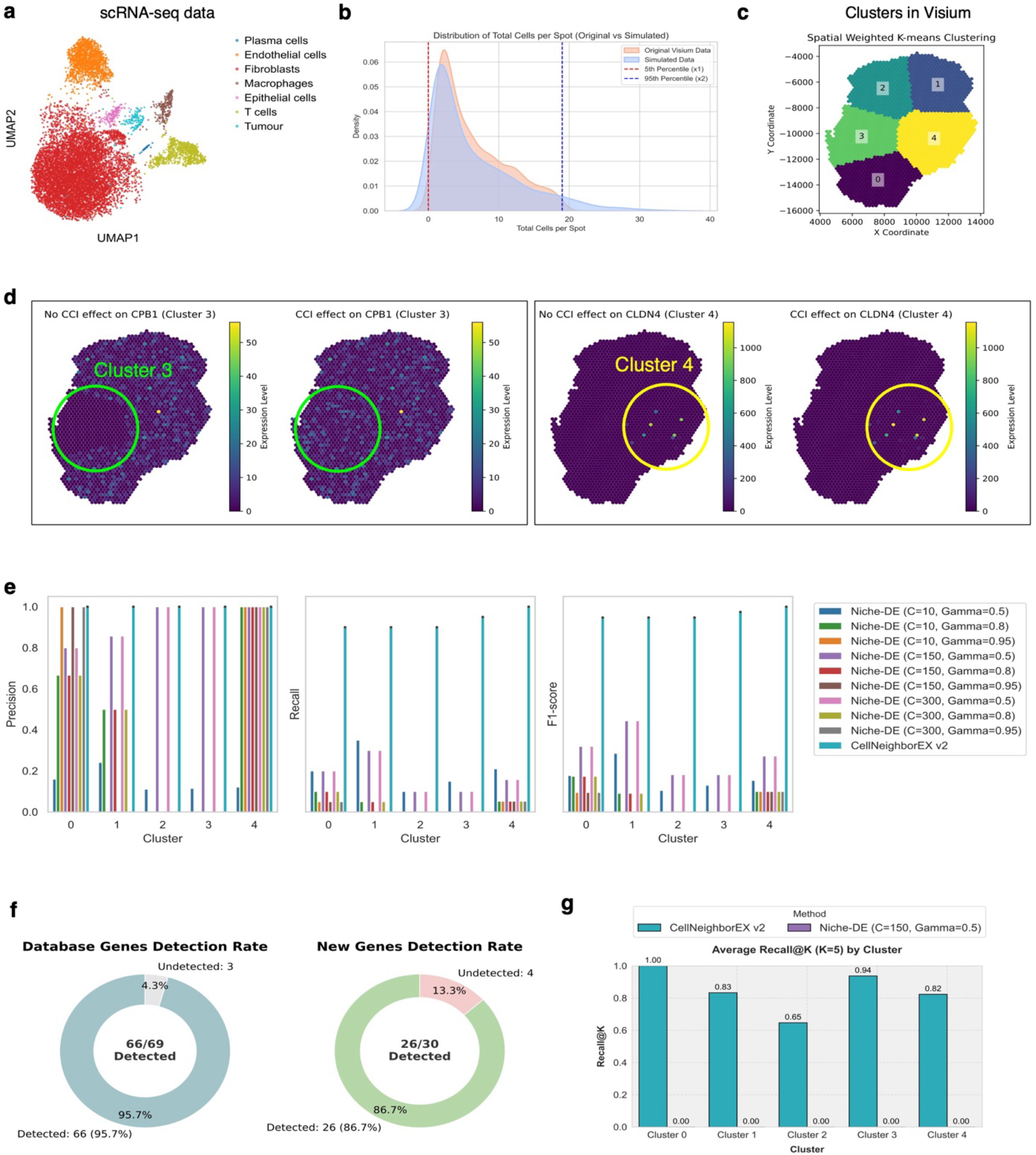
CellNeighborEX v2 accurately recovers context-specific CCI genes in a synthetic spatial transcriptomics benchmark. **a** UMAP visualization of scRNA-seq reference data used to simulate gene expression in the synthetic Visium dataset. **b** Distribution of total cell numbers per spot in the synthetic dataset (orange) closely aligned with the original human ovarian cancer Visium data (blue). **c** Five spatial regions (clusters) defined in the synthetic Visium data, representing distinct biological contexts for evaluating CCI gene detection. **d** Ground-truth CCI gene selection for benchmarking. A total of 99 genes were selected from the scRNA-seq reference dataset, each uniquely associated with one of the five spatial regions. The gene set includes lowly or sparsely expressed genes to create a comprehensive and challenging benchmark for evaluating the performance of CCI detection methods. **e** Performance comparison of CellNeighborEX v2 and Niche-DE across spatial regions using precision, recall, and F1-score. Multiple parameter settings were tested for Niche-DE. **f** Detection rate of database-derived CCI genes (left) and novel, non-database genes (right). CellNeighborEX v2 detects 66 of 69 known genes (95.7%) and 26 of 30 novel genes (86.7%). **g** Recall@K analysis for identifying source-neighbor cell type pairs for each CCI gene.

Cell type proportions for each synthetic spot were sampled from a Dirichlet distribution and scaled by spot-level cell counts to obtain cell type abundances. Gene expression values were then estimated by combining these abundances with scRNA-seq-derived mean expression profiles. Subsequently, spatial clustering identified five distinct regions that served as biological contexts (Fig. 2c, Methods).

To define ground-truth CCI genes, we selected 99 genes uniquely associated with one of the five regions, including lowly or sparsely expressed ones for a challenging benchmark (Fig. 2d). Of these, 69 were ligands, receptors, or signaling components listed in Omnipath^36^, CellChat^37^, or CellTalk^38^, while the remaining 30 were not covered by these ligand-receptor databases. CCI-driven expression was simulated by adding co-abundance-proportional signals with log fold changes between 0.5 and 2.5 (Methods, Supplementary Fig. 2), reflecting biological variation^27,28^.

We applied CellNeighborEX v2 to the synthetic dataset and identified 92 unique spatial region-specific CCI genes (logFC > 0.5, *adjusted p-value* < 0.01), along with their corresponding interacting cell type pairs (Supplementary Data 1 and 2). We benchmarked its performance against Niche-DE^29^ (v0.0.0.9000), another database-independent method designed to detect niche-associated genes from interactions occurring within each Visium spot. Compared with CellNeighborEX v2, Niche-DE directly uses observed expression values without incorporating expected values or accounting for biological context-specific variation. To provide a favorable condition for Niche-DE, we allowed adjustments to its key parameters, including the minimum total expression count per gene (*C*; 10, 150, 300) and the source cell-type expression percentile threshold (*Gamma*; 0.5, 0.8, 0.95), tailoring them to optimize its performance. Additionally, Niche-DE was applied to each spatial region, creating a supportive environment by reducing the search space.

Performance comparison revealed that CellNeighborEX v2 consistently outperformed Niche-DE^29^ across all spatial regions (Fig. 2e). Notably, CellNeighborEX v2 achieved particularly high precision rates, indicating low FP detection, while simultaneously maintaining strong recall. In total, CellNeighborEX v2 detected 66 of 69 database-derived ligand-receptor genes (95.7%) and 26 of 30 non-database genes (86.7%) (Fig. 2f), demonstrating its ability to recover both established and previously uncharacterized interaction-driven signals.

We further evaluated whether the interacting cell type pair for each CCI gene was correctly inferred. For each gene, we ranked all candidate interacting cell type pairs generated by pairing each correlation-filtered source cell type with every other cell type. We then ordered these pairs according to the Wald-test p-value of their regression coefficient and selected the top five (Methods). A prediction was counted as correct if the ground-truth pair appeared within this top five (Recall@5). CellNeighborEX v2 achieved high average Recall@5 across spatial regions, indicating precise attribution of cellular sources (Fig. 2g).

We also applied CellChat^21^ as a representative ligand-receptor-based method to the synthetic dataset. Because CellChat predicts interactions between neighboring spots rather than within individual spots, we applied it to the entire Visium slide instead of individual spatial regions to ensure stable interaction inference. At the gene level, CellChat yielded low precision, recall, and F1 scores (Supplementary Fig. 3a). When assessing the attribution of interacting cell type pairs, CellChat also exhibited very low Recall@5 values (Supplementary Fig. 3b). These results indicate that relying on known databases and inter-spot inference is insufficient to capture context-specific and fine-grained CCI genes from Visium data.

### CellNeighborEX v2 uncovers fine-grained, neighbor-dependent expression in spatial niches

We previously identified CCI genes in high resolution (10 µm) Slide-seq data^14^, which are genes upregulated specifically in spots where two cell types co-localize. Many of these interaction-driven genes exhibited limited overlap with predefined ligand-receptor databases^27^. Since these genes were detected based on spatial proximity within the spot, they serve as a valuable reference for validating short-range interaction predictions. To utilize them for validation purposes, we generated a pseudo-Visium dataset by spatially aggregating Slide-seq spots into 60 µm × 60 µm bins, effectively downsampling the data to approximate Visium-scale resolution (Fig. 3a). This approach allowed us to directly compare interaction detection capabilities between high- and low-resolution platforms using the same underlying tissue sample.

**Fig. 3:**
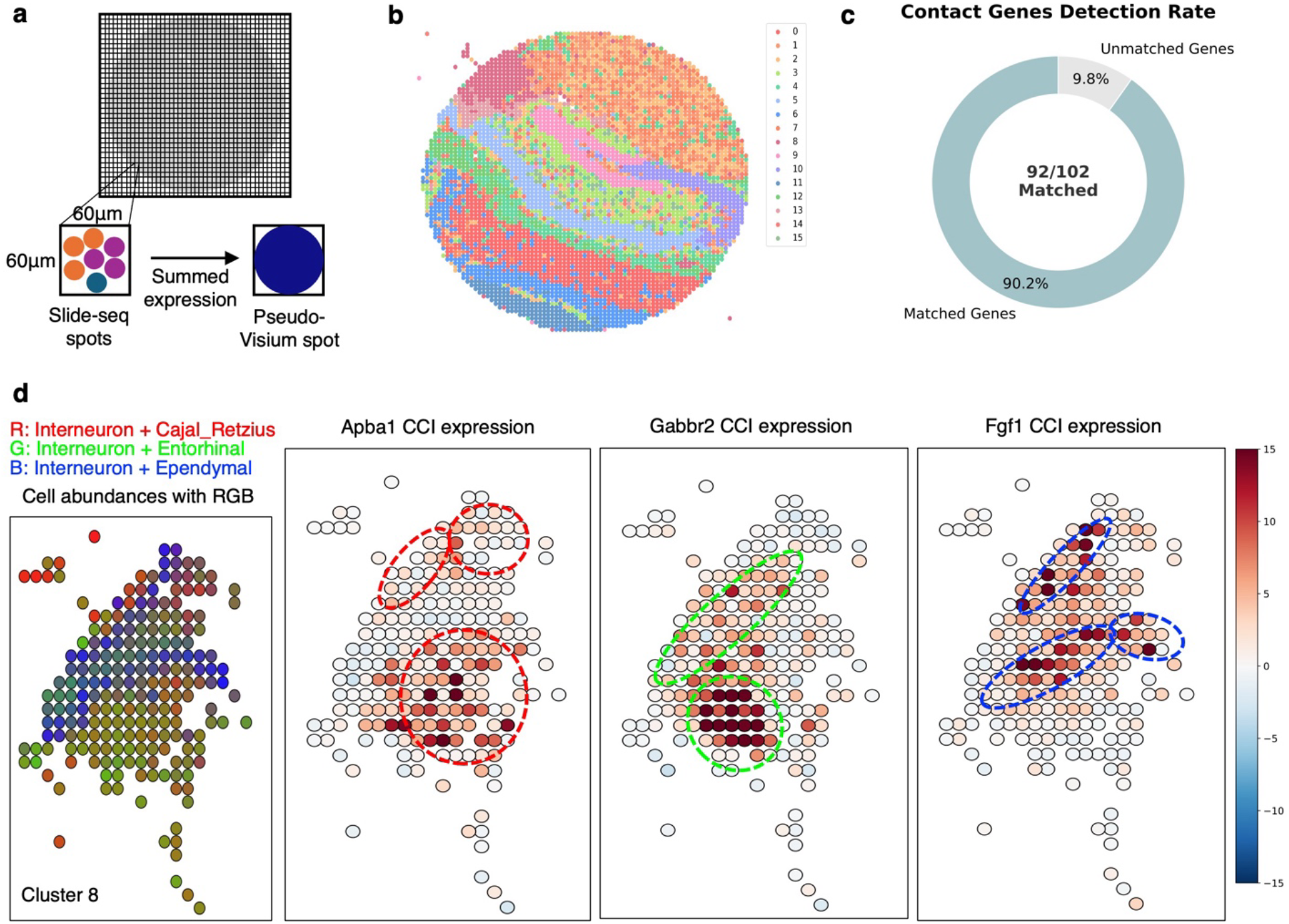
CellNeighborEX v2 uncovers fine-grained, neighbor-dependent CCI gene expression in pseudo-Visium data. **a** A schematic illustration of the generated pseudo-Visium data by aggregating 10 μm Slide-seq spots into 60 μm × 60 μm bins to approximate Visium resolution. **b** Spatial clustering of pseudo-Visium data from the mouse hippocampus defines 16 distinct spatial regions used as biological contexts for CCI gene detection. **c** CellNeighborEX v2 recovered 92 of 102 previously identified cell contact-mediated genes from Slide-seq, demonstrating its ability to extract biologically meaningful signals from low-resolution spatial data. **d** Identified CCI genes correlated with regional cell abundance in spatial region 8 of the pseudo-Visium dataset. Left: RGB visualization of co-abundance patterns for the three predicted pairs (R: interneuron-Cajal-Retzius, G: interneuron-entorhinal, B: interneuron-ependymal). Right: Spatial expression of CCI genes (*Apba1, Gabbr2, Fgf1*) driven by cell-cell interactions.

Spatial clustering of the aggregated spots defined 16 spatial regions (Fig. 3b). Applying CellNeighborEX v2 to the pseudo-Visium data, we identified 1,057 unique spatial region-specific CCI genes (logFC > 0.5, *adjusted p-value* < 0.01) and their corresponding interacting cell type pairs (Supplementary Data 3 and 4). To evaluate sensitivity to short-range interaction signals, we compared our results with 102 genes that we previously identified in the high-resolution Slide-seq data^27^ (Supplementary Data 5). CellNeighborEX v2 successfully recovered 92 of these 102 genes (90.2%) from the downsampled pseudo-Visium data (Fig. 3c), with a highly significant overlap (hypergeometric p-value < 1 × 10^−15^). Of the 10 genes not recovered, five were excluded during scRNA-seq quality control, and the remaining five showed lower observed than expected expression and thus were not classified as CCI genes. This partial loss of recovery results from increased transcript sparsity and signal dilution following spatial downsampling, which obscures subtle interaction signals. Even so, the high overall recovery rate demonstrates that CellNeighborEX v2 can effectively capture fine-grained CCI signals from low resolution Visium data.

To assess whether existing tools capture similar interaction signals, we applied CellChat^21^ to the same pseudo-Visium dataset. Integrating multiple functional categories including cell-cell contact, ECM-receptor, secreted signaling, CellChat identified 318 unique CCI genes. However, it recovered 8 of the 102 previously validated contact-mediated genes, a level of overlap that was not statistically different from random expectation (hypergeometric p-value = 0.989). CellChat models cellular communication primarily at the inter-spot (spot-to-spot) level and is strictly constrained by predefined ligand-receptor databases, making it less sensitive to short-range interactions occurring within individual spots.

Spatial analysis confirmed the colocalization of the predicted interacting cell types of the CCI genes (Supplementary Fig. 4). For instance, in cluster 8, CellNeighborEX v2 identified *Apba1, Gabbr2*, and *Fgf1* as CCI genes, each upregulated through interactions between interneurons and a distinct neighboring cell type: Cajal-Retzius, entorhinal, and ependymal cells, respectively. RGB visualization of these cell type pairs revealed distinct co-abundance zones: red spots indicate enrichment of the interneuron-Cajal-Retzius pair, green for interneuron-entorhinal, and blue for interneuron-ependymal (Methods). Notably, these interaction-specific zones closely matched the CCI expression patterns of *Apba1, Gabbr2*, and *Fgf1* (Fig. 3d). Also, in cluster 13, CellNeighborEX v2 identified *Fabp7* as being expressed in astrocytes when interacting with endothelial tip cells. This interaction in the pseudo-Visium data was spatially consistent with regions where both cell types co-occurred in the matched Slide-seq data (Supplementary Fig. 4).

We extended the analysis to mouse liver cancer Slide-seq^14^ data. Pseudo-Visium data were generated and clustered into nine spatial regions (Supplementary Fig. 5a). CellNeighborEX v2 predicted 1,079 unique spatial region-specific CCI genes (logFC > 0.5, *adjusted p-value* < 0.01) and their corresponding interacting cell type pairs (Supplementary Data 6 and 7). CellNeighborEX v2 recovered 33 of 34 previously identified cell contact-dependent genes (hypergeometric p-value < 6.05 × 10^−10^) (Supplementary Fig. 5b, Supplementary Data 8). Spatial analysis showed that the predicted source and neighboring cell types co-occurred, and their marker genes exhibited spatial co-expression in the matched Slide-seq data (Supplementary Fig. 5c). For instance, *F13a1* was highly expressed in monocytes interacting with tumor cells in a tumor-specific region of the pseudo-Visium data (Supplementary Fig. 5d), resonant to previous observation by original CellNeighborEX^27^.

Our findings confirm that CellNeighborEX can detect short-range interaction-driven gene expression patterns identified from high-resolution Slide-seq data even after spatial downsampling to low-resolution pseudo-Visium data.

### CellNeighborEX v2 characterizes ligand-receptor-mediated communication patterns associated with distinct subclonal microenvironments

To examine the ability of CellNeighborEX v2 in detecting ligand-receptor-mediated CCIs, we analyzed paired Visium and CosMx ST datasets obtained from adjacent sections of a human ovarian cancer sample^35^. Subclonal architecture was defined based on previously annotated malignant clusters, P5.1 and P5.2, identified via copy number alteration (CNA) analysis (Fig. 4a, Supplementary Data 9)^35^. These clusters correspond to spatial regions FOV4 and FOV22, respectively (Fig. 4b). Previous analysis of CosMx data showed that the PIGR+ tumor subclone (FOV4) is surrounded by immune cells, including macrophages and T cells, and expresses immune response genes such as *CXCL10* and *S100A8*^35^. In contrast, the PTGS1+ tumor subclone (FOV22) is enriched with fibroblasts and endothelial cells, and expresses growth factor-related genes such as *BMP7, FGF18*, and *INHBB*^*35*^.

**Fig. 4:**
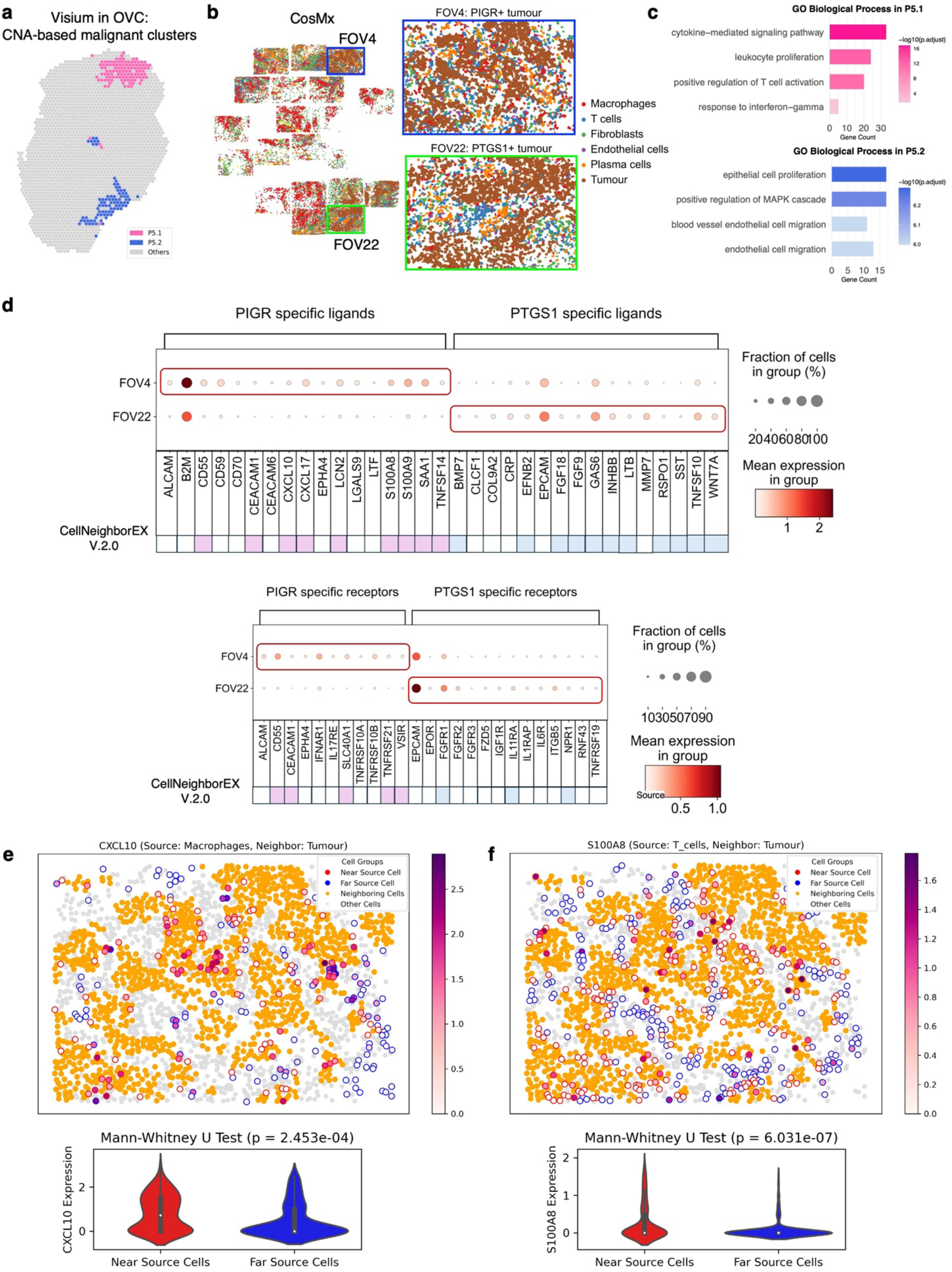
CellNeighborEX v2 reveals tumor subclone-specific microenvironmental heterogeneity in human ovarian cancer. **a** Spatial map of Visium data showing two malignant tumor subclones (P5.1 and P5.2) defined by copy number alteration (CNA) analysis. **b** CosMx high-resolution spatial transcriptomics data from adjacent tissue sections corresponding to P5.1 (FOV4) and P5.2 (FOV22). Tumor cells are shown alongside neighboring immune (e.g., macrophages, T cells) or stromal (e.g., fibroblasts, endothelial) cell types. **c** GO enrichment analysis of CCI genes identified by CellNeighborEX v2 reveals that P5.1 is enriched in immune-related signaling (e.g., cytokine-mediated signaling, T cell activation), whereas P5.2 is enriched in stromal processes (e.g., epithelial proliferation, endothelial migration). **d** Cross-platform validation of CellNeighborEX v2 predictions using CosMx ligand-receptor annotations. Many ligands and receptors enriched in PIGR+ (P5.1) and PTGS1+ (P5.2) tumor subclones in CosMx are also identified as CCI genes in Visium data by CellNeighborEX v2. Dot plots show the mean expression of subclone-specific ligands and receptors in FOV4 and FOV22 across cell types. Overlapped ligands (light pink) and receptors (light blue) were identified both by CellNeighborEX v2 and in the CosMx-defined subclone-specific gene sets. **e** *CXCL10* is predicted to be expressed in macrophages interacting with tumor cells in P5.1. In CosMx data, macrophages near tumor cells exhibit significantly higher expression than those farther away (Mann-Whitney U test p = 2.435e-04). **f** *S100A8* is predicted to be expressed in T cells near tumor cells in P5.1. T cells located closer to tumor cells show higher expression (Mann-Whitney U test p = 6.031e-07), confirming proximity-dependent expression patterns of predicted CCI genes.

Applying CellNeighborEX v2 to the Visium dataset, we identified 213 unique tumor subclone-specific CCI genes (logFC > 0.5, *adjusted p-value* < 0.01) and their associated interacting cell type pairs in P5.1 and P5.2 (Supplementary Data 10 and 11). Gene ontology (GO) enrichment analysis revealed that CCI genes in P5.1 were associated with immune-related processes, including cytokine-mediated signaling and T cell activation. Meanwhile, CCI genes in P5.2 were linked to stromal responses such as epithelial cell proliferation and endothelial cell migration (Fig. 4c). These CCI genes correspond to the biological functions identified in fine-grained CosMx data.

We examined whether CellNeighborEX v2 applied to Visium data could recover known signaling molecules previously identified using the CosMx dataset. Comparing with the curated list of ligands and receptors enriched in the PIGR^+^ and PTGS1^+^ subclone^35^, we found that CellNeighborEX v2 recovered 28 out of 58 ligands and receptors (hypergeometric p-value < 8.57 × 10^−7^) (Fig. 4d).

To confirm the spatial dependency of the interactions predicted by CellNeighborEX v2, we leveraged the CosMx dataset to calculate distances between source and neighboring cell types. For each predicted interaction, source cells were stratified into proximal and distal groups relative to the neighboring cell type, and gene expression was compared. In FOV4, for instance, *CXCL10* was predicted to be expressed in macrophages interacting with tumor cells in P5.1 of the Visium data. Consistent with this prediction, macrophages located near tumor cells exhibited significantly higher *CXCL10* expression than distant ones (Mann-Whitney U Test p-value < 0.05) (Fig. 4e). Similarly, T cells adjacent to tumor cells expressed elevated levels of *S100A8* (Fig. 4f). Multiple additional predicted interactions showed similar spatially dependent expression patterns (Supplementary Fig. 6).

These findings demonstrate that CellNeighborEX v2 can uncover subclone-specific, ligand-receptor-driven intercellular communication and accurately resolve microenvironmental heterogeneity from low-resolution Visium data.

### CellNeighborEX v2 identifies a tumor-TME crosstalk hotspot and its associated functional CCI genes

To measure the ability of CellNeighborEX v2 to identify functional CCI genes associated with tumor-TME interactions, we analyzed paired Visium and Visium HD datasets derived from the same human colorectal cancer tissue section^12^. Seven spatial regions were obtained from the Visium data (Supplementary Fig. 7). CellNeighborEX v2 detected 3,317 unique spatial region-specific CCI genes (logFC > 0.5, *adjusted p-value* < 0.01) and their source and neighboring cell type pairs (Supplementary Data 12 and 13).

Spatial mapping to high-resolution data confirmed that the predicted cell types were indeed co-localized in Visium HD, and that the CCI genes were highly expressed in those regions (Supplementary Fig. 8 and 9). For a comparative analysis, we applied CellChat^21^, which performs spot-to-spot analysis, to the Visium HD data (Methods). Among the gene-source-neighboring cell type triplets identified by CellChat, CellNeighborEX v2 recovered 9 signaling-, 35 ECM-related, and 14 contact-mediated genes. For instance, *SPP1* (signaling), *COL1A2* (ECM), and *CD177* (contact) showed consistent spatial expression in the predicted interacting cell types (hypergeometric p-value < 1 × 10^−15^) (Fig. 5a-c, Supplementary Fig. 10-12). Importantly, while CellChat identified these genes in high-resolution Visium HD, it failed to detect them in Visium data. This demonstrates the superior capability of CellNeighborEX v2 in capturing CCI signals even at lower resolutions.

**Fig. 5:**
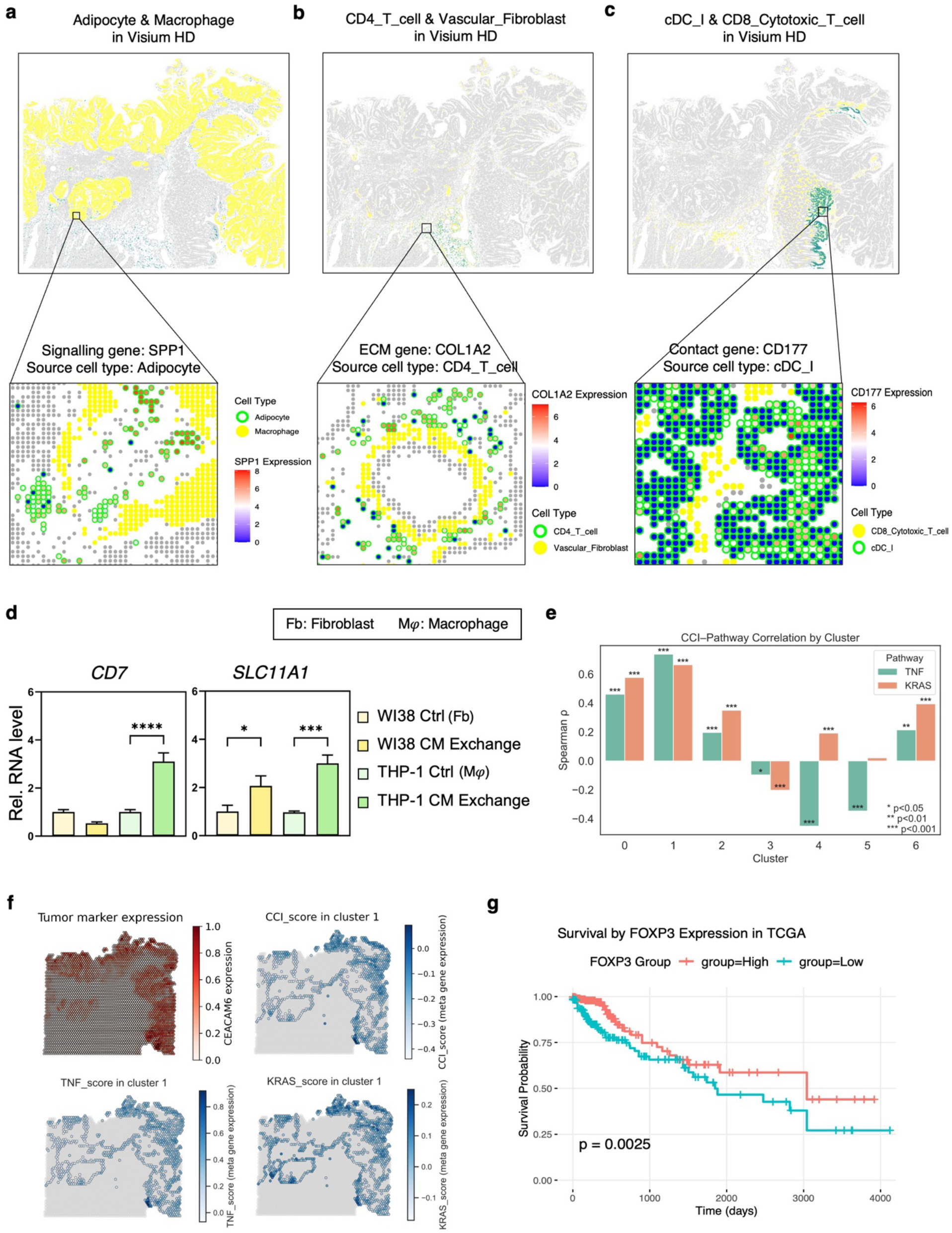
CellNeighborEX v2 identifies a tumor-microenvironment crosstalk hotspot and functionally relevant CCI genes in colorectal cancer. **a-c** Spatial validation of CellNeighborEX v2-predicted CCI genes and interacting cell type pairs using high-resolution Visium HD data. CellNeighborEX v2 was applied to Visium data to identify spatial region-specific CCI genes and predict source-neighboring cell type pairs. The corresponding Visium HD dataset was used to confirm the co-localization of the predicted cell types and the expression of CCI genes at higher resolution. Genes shown here were also recovered by CellChat applied to the Visium HD data and represent distinct modes of cell-cell communication: (a) *SPP1* (paracrine signaling), expressed in adipocytes interacting with macrophages, (b) *COL1A2* (ECM remodeling), expressed in CD4^+^ T cells interacting with vascular fibroblasts, and (c) *CD177* (cell-cell contact), expressed in cDC1 interacting with CD8^+^ cytotoxic T cells. **d** Validation of CellNeighborEX v2-specific CCI genes *CD7* and *SLC11A1* using conditioned medium (CM) experiments. THP-1-derived macrophages cultured in fibroblast-derived CM exhibited significantly increased expression of *CD7* and *SLC11A1*, supporting interaction-dependent induction. Quantitative RT-PCR analysis confirmed that *CD7* expression was specific to macrophages, while both genes were upregulated upon exposure to fibroblast-derived signals. Data are represented as mean ± SE (n = 3); statistical significance was assessed using one-way ANOVA. **e** Correlation between meta-gene scores for TNF-α and KRAS pathway genes and those of CellNeighborEX v2-predicted CCI genes across Visium-defined spatial regions. Cluster 1 exhibits the strongest positive correlation with both pathways. **f** Spatial co-localization of tumor marker *CEACAM6*, TNF-α and KRAS pathway meta-gene scores, and a meta-gene score representing enriched CCI genes in cluster 1 of the Visium data. **g** Kaplan-Meier survival analysis shows that high expression of *FOXP3*, a known immune suppressive regulator, is associated with improved survival in TCGA colorectal cancer patients.

We further experimentally validated long-range CCI genes predicted by CellNeighborEX v2 but not by CellChat when applied on the Visium HD data. Among them, *CD7* and *SLC11A1* were predicted to be upregulated in macrophages interacting with fibroblasts (Wald-test p-value = 0.0045 and p-value = 0.0233, respectively). To test this, macrophages were cultured in media previously used to culture fibroblasts (Methods). Both *CD7* and *SLC11A1* showed increased expression in macrophages exposed to fibroblast-derived conditioned media (CM) (Fig. 5d). Similar results were confirmed in transwell experiments (Supplementary Fig. 13).

CellNeighborEX v2 identified genes associated with two major oncogenic pathways involved in tumor-TME crosstalk in colorectal cancer: TNF-α signaling via NF-κB and KRAS signaling^39-43^. Gene sets corresponding to these pathways were obtained from the MSigDB database (Supplementary Data 14 and 15), and their transcriptional activity was summarized as meta-gene expression scores (Methods). CCI genes in cluster 1 exhibited the strongest correlations with both TNF-α and KRAS pathway scores (Fig. 5e) and displayed significant gene overlap (Supplementary Fig. 14). Also, *CEACAM6*^*12*^, a known tumor marker, was highly expressed in cluster 1, co-localizing with increased TNF-α and KRAS scores and enriched CCI gene expression (Fig. 5f).

CellNeighborEX v2 also detected *FOXP3*, a known master regulator of immune suppression in the TME^44,45^. Analysis of TCGA cancer samples revealed that high *FOXP3* expression is significantly associated with improved patient survival, supporting its functional relevance (Fig. 5g) (Methods). Consistently, multiple studies in colorectal cancer cohorts have shown that high expression of *FOXP3* is associated with favorable clinical outcomes, including improved overall and disease-free survival^46-48^. These findings indicate that CellNeighborEX v2 can effectively capture fine scale microenvironmental interactions and identify functionally relevant CCI genes. Notably, *FOXP3* was not detected by CellChat due to its low expression level in Visium HD. Over 99.9**%** of Visium HD spots exhibited zero *FOXP3* transcripts (Supplementary Fig. 15).

### CellNeighborEX v2 reveals condition-specific CCI genes overlooked in single-cell analyses

To investigate whether CellNeighborEX v2 can identify condition-specific CCIs genes, we analyzed paired Visium and scRNA-seq datasets derived from lymph nodes of Mycobacteria-infected (MS) and control (PBS) mice^49^. We leveraged previously processed and batch-corrected scRNA-seq data (Fig. 6a), annotated with six major immune cell types (Fig. 6b)^49^. Corresponding Visium datasets were available from the same biological conditions (Fig. 6c).

**Fig. 6:**
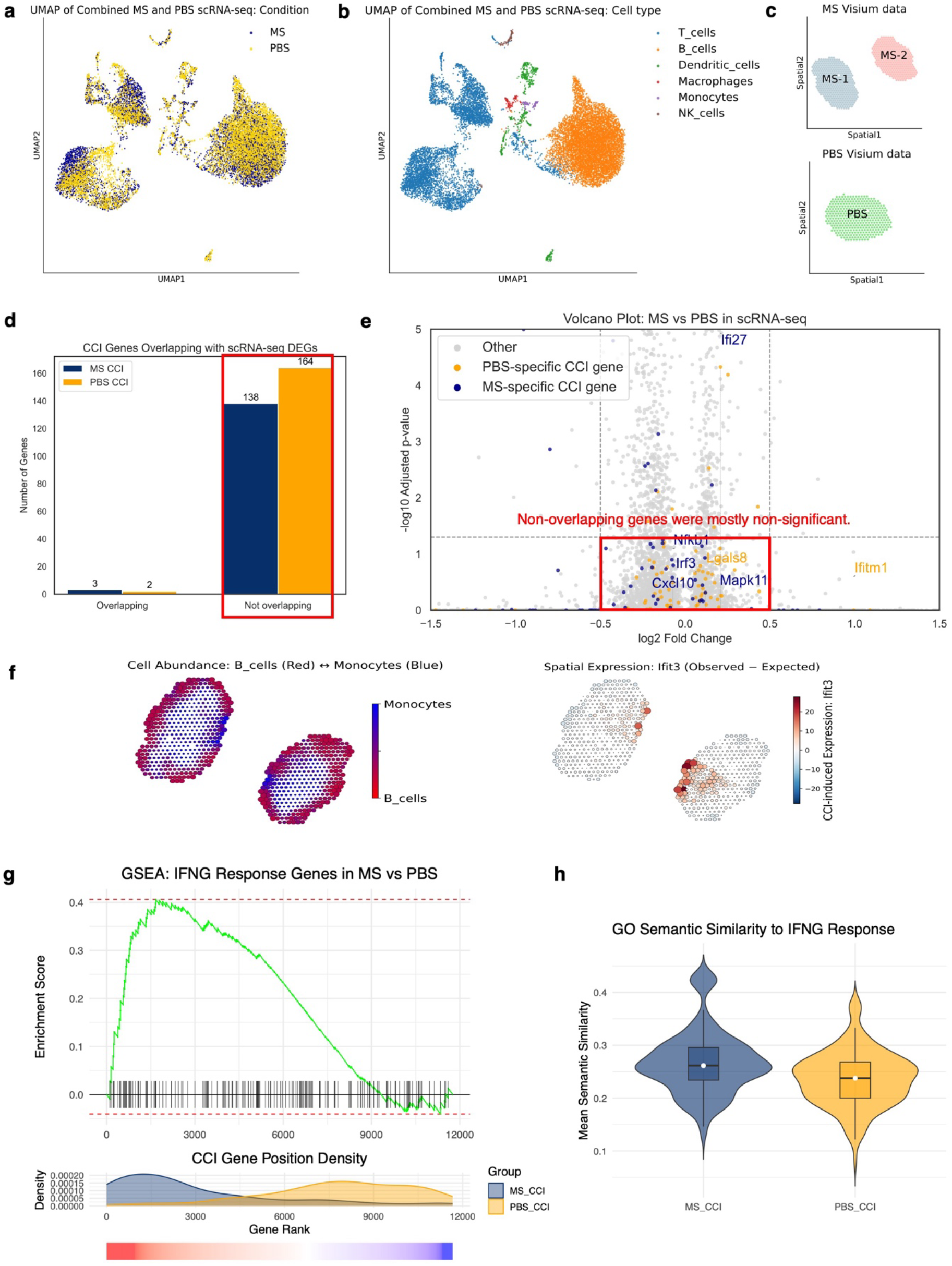
CellNeighborEX v2 identifies IFN-γ-associated, condition-specific CCI genes not captured by single-cell DEG analysis. **a** UMAP visualization of scRNA-seq data from control (PBS) and Mycobacteria-infected (MS) mouse lymph nodes. **b** Annotated scRNA-seq cell types used for spatial deconvolution. **c** Spatial transcriptomic data from matched Visium slides for PBS and MS conditions. **d** Bar plot showing minimal overlap between condition-specific CCI genes identified by CellNeighborEX v2 and DEGs from scRNA-seq analysis using Wilcoxon test. **e** Volcano plot showing that most condition-specific CCI genes were not statistically significant in the single-cell DEG analysis. **f** Spatial expression of *Ifit3*, an interferon-inducible gene, driven by cell-cell interactions, is specifically enriched in regions of monocyte-B cell interaction under the MS condition. **g** Gene set enrichment analysis (GSEA) of Visium-ranked genes reveals significant enrichment of IFN-γ response genes in the MS condition. MS-specific CCI genes are enriched near the top of the list, while PBS-specific genes are biased toward the lower ranks. **h** Semantic similarity scores between gene ontology (GO) terms associated with each CCI gene and those of the IFN-γ response gene set. MS-specific CCI genes exhibit significantly higher similarity (Wilcoxon test, ***p < 0.001).

Applying CellNeighborEX v2 to the Visium datasets, we identified 328 unique experimental condition-specific CCI genes (logFC > 1.0, *adjusted p-value* < 0.01) and their corresponding interacting cell type pairs (Supplementary Data 16 and 17). To facilitate direct comparison between the two conditions, we excluded genes present in both conditions and obtained mutually exclusive CCI gene sets (Supplementary Fig. 16). We then investigated whether the condition-specific CCI genes could also be identified through differential expression analysis of scRNA-seq data. We performed the Wilcoxon rank sum test to obtain differentially expressed genes (DEGs) for each condition (logFC > 0.5, *adjusted p-value* < 0.05) (Supplementary Data 18). Notably, only 5 of the condition-specific CCI genes overlapped with scRNA-seq DEGs (Fig. 6d). This limited overlap was primarily due to most CCI genes not reaching statistical significance or sufficient fold changes in the DEG analysis (Fig. 6e).

Among MS-specific CCI genes, *Ifit3* is a well-known interferon-induced gene involved in immune activation^50,51^. Our analysis showed that *Ifit3* was highly expressed in regions of monocyte-B cell interaction in the MS Visium data (Fig. 6f). This pattern was also experimentally validated by previous immunofluorescence imaging studies in infected lymph nodes^49^, suggesting that *Ifit3* can be induced by interactions between monocytes and B cells. Importantly, *Ifit3* was not discovered from scRNA-seq analysis to be MS-specific. ScRNA-seq DEG analysis even identified it as PBS-upregulated.

Given that MS samples represent an infection model, we examined whether the MS-specific CCI genes were enriched for interferon gamma (IFN-γ) response, a key pathway activated during infection^52,53^. The IFN-γ response gene set was retrieved from MSigDB (Supplementary Data 19). We performed gene set enrichment analysis (GSEA) using a gene list ranked by log fold changes from Visium data (Methods). We confirmed significant upregulation of IFN-γ response genes in the MS condition (Fig. 6g). When we projected the positions of CCI genes onto the ranked list, MS-specific CCI genes were concentrated toward the MS-upregulated region, whereas PBS-specific genes skewed toward the PBS end. This positional bias suggests that MS-enriched CCI genes are functionally linked to IFN-γ signaling.

To further quantify this relationship, we computed the semantic similarity between GO terms associated with each CCI gene and those related to the IFN-γ response gene set (Methods). MS-specific CCI genes exhibited significantly higher similarity scores compared to PBS-specific ones, supporting their relevance to IFN-γ immune processes (Fig. 6h).

## Discussion

The development of NGS-based ST technologies has steadily progressed toward platforms with ever higher spatial resolution, raising expectations for improved detection of CCIs. In principle, such high-resolution approaches should offer unparalleled insights into cellular and subcellular features. In practice, however, they are hindered by substantially reduced sensitivity, particularly for low-abundance transcripts^11,13^. This limitation often necessitates spatial aggregation or binning of high-resolution data to near-Visium resolution to achieve sufficient signal, effectively negating the benefits of increased resolution^16-20^. Consequently, the move toward higher resolution has not yet translated into clear advances in CCI analysis. Additionally, image-based ST platforms are limited by restricted gene panels and a potential increase in technical noise as panel sizes expand, hindering their effectiveness for the comprehensive discovery of unknown CCI genes.

Visium can be a practical alternative to study CCI. Despite its relatively low resolution, Visium has become as the most widely adopted ST platform across diverse tissue types and disease contexts. It offers comprehensive transcriptome coverage and strong compatibility with diverse biological samples^10^. However, beyond ligand-receptor analysis, studying CCI with Visium data remains challenging. This is mainly due to difficulties in accurately locating individual cells, which prevents a comprehensive analysis of diverse communication modalities. A common simple practice is to annotate each Visium spot with its dominant cell type, as implemented in tools like CellChat^21^ and NCEM^25^. These methods primarily assess interactions between neighboring spots, overlooking the complex multicellular interactions that occur within individual spots. As a result, they are unable to capture the full spectrum of intercellular communication patterns present in tissue microenvironments. While Niche-DE^29^ inferred interactions from Visium data without the reference, its practical utility is highly limited by a high rate of FP predictions (Fig. 2).

CellNeighborEX v2 offers a new framework for investigating multiple modes of CCIs using Visium data. Even at the relatively low resolution of Visium, it can detect both short- and long-range interactions. The framework models deviation of the observed expression relative to the expected values derived from matched scRNA-seq data and cell type deconvolution. This residual-based approach works independently of predefined ligand-receptor databases, enabling discovery of non-canonical and novel CCI genes across diverse communication modalities.

CellNeighborEX v2 identifies candidate CCIs by quantifying the deviation of observed gene expression from expected values. We recognize that gene expression deviations can arise from technical noise or intrinsic biological gradients (*e*.*g*., metabolic or developmental zonation) unrelated to cellular proximity. To mitigate these confounding effects, our permutation framework preserves local cell type composition and spatial context while randomizing neighbor relationships, thereby testing whether observed residual signals depend specifically on local cell type co-occurrence rather than positional gradients alone.

The framework is motivated by observations that genes involved in intercellular communication are often upregulated relative to other cells of the same cell type, as revealed by physically interacting cells and high-resolution ST data^27,28^. Consistent with this, analyses of scRNA-seq data have shown that only subsets of cells within a given cell type express specific ligands or receptors, and these expressing cells typically exhibit higher expression levels compared to the average expression of that gene within the same cell type as used by CellPhoneDB^54^ and NicheNet^55^.

Extensive validation across synthetic and biological datasets confirms the accuracy of CellNeighborEX v2 in predicting CCI genes and their cellular sources. Compared with Niche-DE, CellNeighborEX v2 achieved substantially higher performance, highlighting the effectiveness of our statistical tests in reducing false positives and detecting diverse modes of CCIs (Fig. 2). The framework successfully recovered CCI genes from pseudo-Visium data that were previously identified in the original high-resolution Slide-seq datasets (Fig. 3). While benchmarking with real Visium datasets is challenging due to the lack of exhaustive ground-truth knowledge, CellNeighborEX v2 consistently recapitulated interactions observed in matched high-resolution CosMx or Visium HD data from adjacent sections (Fig. 4 and Fig. 5). These results, including those from the pseudo-Visium analysis, demonstrate that CellNeighborEX v2 can uncover key CCI signals that are otherwise undetectable using conventional approaches (Fig. 3).

Moreover, CellNeighborEX v2 detected CCI genes such as *FOXP3* from Visium data even in cases where higher-resolution platforms such as Visium HD fail due to limited transcript detection sensitivity (Fig. 5g, Supplementary Fig. 15). CellNeighborEX v2 also identified macrophage-specific expression changes driven by long-range CCIs, which was not detected using ligand-receptor based CellChat^21^. The induction of these genes was further validated using CM assays (Fig. 5d). Besides, it revealed CCI changes that are undetectable by scRNA-seq CCI analysis, underscoring the critical importance of spatial context in resolving intercellular communication events masked in dissociated single-cell data (Fig. 6).

Another key strength of CellNeighborEX v2 is its ability to capture context-specific CCIs, identifying transcriptional changes in cellular communication across different biological contexts such as spatial regions, disease subtypes, and experimental conditions. In mouse hippocampus pseudo-Visium data, the method revealed spatial region-specific, neighbor-dependent gene expression (Fig. 3). In human ovarian cancer, it resolved tumor subclone-specific interactions associated with distinct microenvironmental programs (Fig. 4). In colorectal cancer, it identified a spatial tumor-microenvironment interaction hotspot involving TNF-α and KRAS signaling pathways (Fig. 5). In the mouse lymph node, it discovered immune response genes such as *Ifit3* from the Mycobacteria-infected Visium sample (Fig. 6). These diverse applications demonstrate that CellNeighborEX v2 not only reliably detects CCIs but also provides critical insights into the dynamic and context-dependent nature of intercellular communication, making it a powerful tool for advancing our understanding of complex tissue microenvironments.

Although this study primarily utilizes Visium, the statistical framework of CellNeighborEX v2 is inherently platform-agnostic and applicable to any multicellular spot-based ST data. Our successful validation using spatially aggregated Slide-seq data, which shares NGS-based characteristics and transcript sparsity with platforms such as DBiT-seq^9^ and Stereo-seq^16,17^, underscores the broad extensibility of our method. While high sparsity in certain technologies may pose challenges for deconvolution, CellNeighborEX v2 effectively mitigates these issues by leveraging residual-based modeling to capture interaction signals that persist despite transcript dilution.

CellNeighborEX v2 supports comprehensive characterization of intercellular communication in large-scale Visium datasets, advancing applications such as biomarker discovery, immune profiling, and spatial disease monitoring. By uncovering both established and previously unrecognized interactions, CellNeighborEX v2 addresses key limitations of existing methods and broadens the analytical scope of spatial transcriptomics. With the abundance of publicly available Visium datasets, CellNeighborEX v2 provides a powerful framework to reanalyze these resources and uncover deeper insights into CCI mechanisms, including those shaping the tumor microenvironment and developmental trajectories.

## Methods

### Synthetic data generation

To evaluate the ability of CellNeighborEX v2 to recover context-specific CCI genes, we generated a synthetic ST dataset designed to mimic the spatial and biological complexity of real Visium data. Spot coordinates and the number of cells per spot were extracted from a human ovarian cancer Visium dataset^35^ to preserve realistic spatial architecture and cell types.

We used matched, annotated scRNA-seq data^35^ to estimate spot-level cell abundance. Cell type deconvolution was performed on real Visium data using cell2location^56^ (v0.1.3). To capture biologically plausible variation, cell counts for each synthetic Visium spot were sampled within the 5th to 95th percentile range of the cell count distribution derived from real data. To assign specific cell type proportions and abundances in each synthetic spot, we sampled from a Dirichlet distribution and multiplied the proportions by the total cell count per spot to obtain cell type abundances.

For computational efficiency during repeated simulations, the total number of genes was set to approximately 650. The corresponding reference gene expression signatures for each cell type were estimated using a negative binomial regression model implemented in the cell2location framework^56^, which models gene expression while accounting for overdispersion and technical noise. The resulting signatures represent the average expression levels per gene for each annotated cell type.

Synthetic gene expression values were computed by multiplying the per-spot cell type abundances with the corresponding reference gene expression signatures. To simulate spatial contexts, we performed K-means clustering with k=5 using features that combine spatial coordinates and spatially weighted gene expression profiles. Specifically, we computed spatially weighted expression by applying a Gaussian kernel to pairwise spatial distances between spots. The original expression matrix was then combined with the spatially smoothed matrix using a spatial weight parameter (λ = 0.5).

As a negative control, we first applied CellNeighborEX v2 to the synthetic dataset without CCI-driven effects and verified the absence of false positives. Subsequently, to define ground truth CCI genes, we selected 99 genes across the five clusters: 69 known ligand-receptor or interaction-related genes from Omnipath^36^, CellChat^37^, or CellTalk^38^ databases, and 30 non-database genes. The selected genes were mutually exclusive across clusters. Each selected gene was assigned a pair of interacting cell types (source and neighboring), chosen from the dominant cell types in each spatial region. The “source” cell type refers to the cell type in which the CCI gene is expressed, whereas the “neighboring” cell type is its spatially adjacent partner assumed to interact with the source.

To reflect the variance typically observed in real datasets, we added noise to the expression values of the ground truth genes. For each gene, a dispersion parameter was estimated from the scRNA-seq data under a negative binomial distribution assumption and applied to the synthetic matrix to simulate biological overdispersion.

We added synthetic CCI-driven expression signals to these ground truth genes to reflect spatially localized cell-cell communication. The interaction strength per spot was defined as the product of source and neighboring cell type proportions:

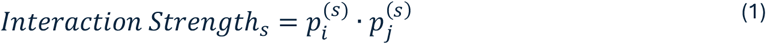

Only spots with interaction strength above the 5th percentile were selected for signal injection. The CCI expression signal added to each gene *g* in spot *s* was defined as:

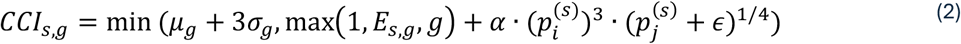

where:

- *μ*_*g*_ and *σ*_*g*_ are the mean and standard deviation of gene *g*’s expression across all spots,
- *E*_*s, g*_ is the original expression of gene *g* in spot *s*,
- *α* is a scaling constant (set to 500),
- *ϵ* is a small constant (set to 10^-5^) to avoid division by zero.

This formulation places higher weight on the source cell type and modulates the neighboring cell contribution more gently, capturing directional characteristics of cell-cell signaling. This design reflects the expectation that CCI gene expression is typically stronger in source cells compared to neighboring cells. All resulting expression values were capped at *μ*_*g*_ + 3*σ*_*g*_ to prevent unrealistic overexpression.

We validated the quality of synthetic data by reapplying cell2location^56^ to predict cell type abundance and proportions from the synthetic expression matrix. These predictions were compared to the original ground truth values derived from the Dirichlet distribution. Specifically, mean squared error (MSE) and Pearson correlation were calculated. These metrics showed strong consistency (Supplementary Data 20 and 21), confirming that the deconvolution framework accurately recovered the simulated cell type compositions.

This synthetic dataset provides a controlled and biologically realistic benchmark for evaluating the performance of CellNeighborEX v2 in recovering spatial region-specific CCI genes. Furthermore, it offers a reproducible testbed for comparative analyses and methodological development in low-resolution spatial transcriptomics.

### Estimation of expected expression

To identify CCI-driven gene expression, we first estimated the baseline expression of each gene *g* at each Visium spot *s*, referred to as the expected expression. This value represents the gene expression predicted solely based on the cell type composition at each spot.

Two key components were used as inputs:

- *ρ*_*f,g*_: the average expression of gene *g* in cell type *f*, estimated from raw count data in annotated scRNA-seq using a negative binomial regression model,
- *α*_*s,f*_: the abundance of cell type *f* in spot *s*, inferred through deconvolution of Visium data.

To model additional biological and technical confounders, we used the cell2location framework^56^:

- *d*_*g*_: gene-specific scaling factors capturing sensitivity differences between scRNA-seq and spatial platforms,
- *b*_*e,g*_: batch- or slide-specific biases for gene *g* in batch or slide *e*,
- *l*_*s*_: latent effects associated with the spatial location of spot *s*.

Thus, the expected expression for gene *g* at spot *s* is then given by:

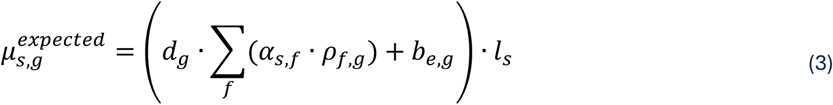

All parameters were estimated using variational Bayesian inference^56^, which fits a generative model of transcript count data conditioned on reference expression signatures and spatial cell type composition.

Finally, we computed residuals by subtracting the expected expression from the observed gene expression counts 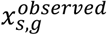 measured in Visium:

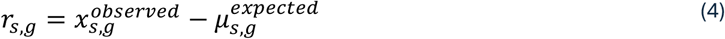

These residuals reflect the extent to which observed gene expression exceeds expectation and thus may serve as indicators of CCI-driven transcriptional activity.

### Hybrid statistical framework for CCI gene detection

To identify context-specific CCI genes, we devised a hybrid statistical framework that combines a goodness-of-fit test within biological contexts and a permutation-based comparison across contexts. Here, a biological context is defined as a spatial region. For each gene, we assessed whether its CCI-driven expression was significantly enriched within a given context.

#### (i) Chi-squared test within a context

To detect genes whose expression significantly deviates from expectation within a given context, we applied a chi-squared (χ^2^) goodness-of-fit test to compare observed versus expected gene expression across spots in each cluster. The test statistic for gene *g* in context *c* is defined as:

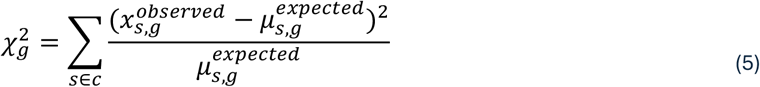

where 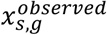 is the observed expression and 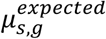 is the expected expression at spot *s*. To ensure statistical robustness, we excluded spots with zero observed expression (i.e., use_zero=False), since genes inactive in a region are not expected to contribute to CCI-specific signals. This strategy was also applied when calculating log fold-change (logFC). Multiple testing correction was performed using the Benjamini-Hochberg method to obtain adjusted p-values.

#### (ii) Permutation test between contexts

To further ensure that identified genes are specifically upregulated in one context relative to others, we performed a permutation test across contexts. For each gene, residual expression values 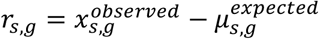 were randomly shuffled across spots 1,000 times. For each permutation *i*, we computed the mean residual for gene *g* in context *k*, denoted 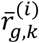.

The empirical null distribution was then used to compute a z-score for the observed mean residual 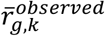 , which was converted to a p-value using the cumulative distribution function (CDF) of the standard normal distribution:

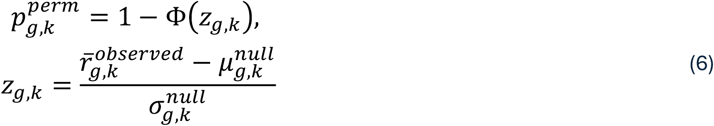

Here, Φ(·) denotes the standard normal CDF. For the ground truth CCI genes in the synthetic dataset, permutation-based null distributions largely passed the Shapiro-Wilk normality test (Supplementary Fig. 17), supporting the validity of Z-score-based inference. Only a negligible proportion of gene-cluster combinations (<10%) showed significant deviation from normality. This Z-score-based approach provides a smooth, continuous p-value distribution, which is required for combination with the chi-squared p-values via the Cauchy method to ensure numerical stability. The p-values were also adjusted using the Benjamini-Hochberg procedure.

#### (iii) Cauchy combination of p-values

In synthetic data experiments, we independently evaluated the performance of each test, and found that the permutation test reduced false positives, while the chi-squared test mitigated false negatives (logFC > 0.5, *adjusted p-value* < 0.01) (Supplementary Fig. 18). To leverage the complementary strengths of both tests, we applied the Cauchy combination test^33,34^ to integrate their p-values:

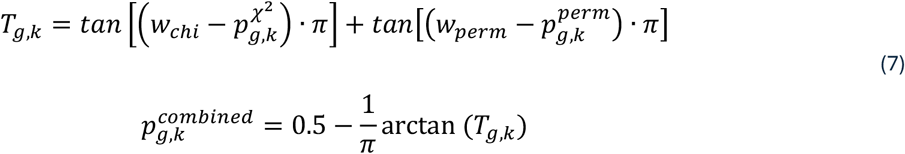

To determine the optimal weighting scheme, we systematically varied the chi-squared (*w*_*chi*_) to permutation weight (*w*_*perm*_) ratios from 0.1 to 0.9 and evaluated the detection accuracy using synthetic datasets. We defined sensitivity as the proportion of ground truth gene-cluster pairs detected with *adjusted p-value* < 0.01, and specificity as one minus the false positive rate among non-ground truth pairs. The harmonic mean of sensitivity and specificity was used to score each combination. A 0.9:0.1 weighting of chi-squared and permutation p-values yielded the highest score (Supplementary Fig. 19), indicating the most favorable balance between sensitivity and specificity.

### Two-step regression for identifying interacting cell type pairs

To identify the source and neighboring cell type pairs for each context-specific CCI gene, we implemented a two-step regression framework. This strategy adjusts for confounding influences due to the complex cellular composition of each spot and allows quantification of interaction-specific effects.

#### (i) Candidate source cell type selection

For each context-specific CCI gene *g*, we first computed scaled residual expression values to identify candidate source cell types. The scaled residual at spot *s* was calculated by subtracting the expected expression 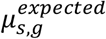 from the observed expression 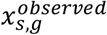, and dividing by the square root of the expected expression to stabilize variance:

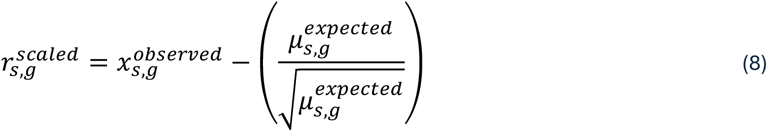

This scaling was applied to mitigate the proportional increase of both expected and observed expression values, which arises from the negative binomial distribution and the linearity of the model estimating expected expression.

To attribute gene expression to specific cell types, we computed the cell type-specific contribution at each spot *s* as:

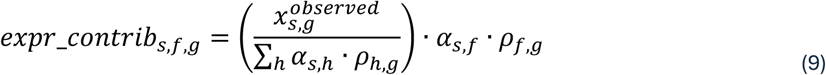

Here, *α*_*s,f*_ denotes the deconvolved abundance of cell type *f* at spot *s*, and *ρ*_*f,g*_ is the average expression of gene *g* in cell type *f*, inferred from scRNA-seq data. The denominator Σ_*h*_ *α*_*s,h*_ · *ρ*_*h,g*_ estimates the total expected contribution of all cell types to the observed expression of gene *g* at spot *s*. This formulation proportionally distributes the observed expression 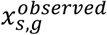 across cell types according to their local abundance and expression signatures, resulting in an estimated contribution of each cell type to gene expression per spot.

We then assessed the Spearman correlation between the scaled residuals 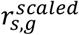 and the cell type specific contributions *expr_contrib*_*s,f,g*_ across all spots in a given context. Cell types with correlation ≥ 0.6 and *p-value* < 0.05 were selected as candidate source cell types for further regression modeling. Cell types exhibiting strong positive correlation were considered primary contributors to the residual expression of the gene, potentially indicating CCI-driven effects.

#### (iii) Interaction modeling with two-step regression

To quantify the influence of neighboring cell types on the expression of CCI genes, we modeled residuals 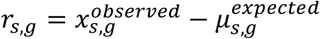 using a regression framework. In contrast to the source cell type selection step, we used unscaled residuals here because they directly represent the magnitude of unexplained expression beyond expectation and thus offer a more interpretable measure of effect size. The validity of this approach is supported by the observed overdispersion of residuals for ground truth genes relative to the Poisson reference line (Supplementary Fig. 20).

We defined interaction terms between a candidate source cell type *f*_*o*_ and each neighboring cell type *f*_*n*_ at each spot *s* as:

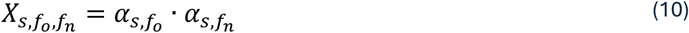

We first fit a full non-negative negative binomial regression model with ridge regularization:

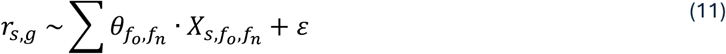

where 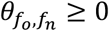 are the regression coefficients indicating interaction effects between source *f*_*o*_ and neighbor *f*_*n*_, and *ε* is the noise term. Because our analysis focused on CCI genes that are upregulated beyond expected expression levels, we imposed a non-negativity constraint on the interaction coefficients. Ridge regularization controls for multicollinearity by penalizing the squared magnitude of coefficients. To minimize potential confounding from self-interactions, we restricted all interaction modeling to heterotypic cell type pairs (*f*_*o*_ ≠ *f*_*n*_). Although our analyses focused on heterotypic interactions, self-interaction terms were implemented in the modeling framework and can be optionally enabled.

To isolate the marginal contribution of a specific source-neighbor pair, we conducted a second regression retaining only the focal interaction term as the independent variable and treating the estimated effects of other terms as an offset:

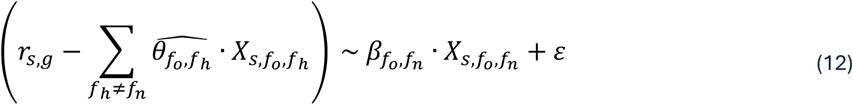

This two-step approach enabled us to distinguish the unique effect of a given cell type pair while adjusting for the confounding influence of other interactions.

To statistically evaluate the contribution of each interaction term, we assessed the significance of each estimated coefficient 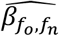 by computing the corresponding Wald test p-value:

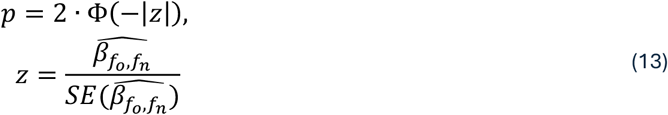

where 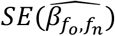 is the standard error of the estimated coefficient and Φ is the cumulative distribution function of the standard normal distribution. These p-values were used to rank candidate interacting cell type pairs for each CCI gene.

#### (iii) Classification of novel CCI genes via database comparison

To identify potentially novel CCI genes not documented in existing interaction resources, we systematically compared our predicted gene list against curated ligand-receptor databases. Specifically, we used OmniPath^36^, CellChatDB^37^, and CellTalkDB^38^, which compile experimentally validated and computationally inferred ligand-receptor and source-target gene interactions in human and mouse.

Any gene not found in these reference databases was classified as a novel CCI gene. This classification allowed us to distinguish between previously known interaction-related genes from newly inferred candidates, which may serve as promising targets for future experimental validation and mechanistic investigation.

### Biological datasets and preprocessing for CellNeighborEX v2 applications

We applied CellNeighborEX v2 to five ST datasets across mouse and human tissues to identify and validate context-specific CCI genes. In all cases, we performed cell type deconvolution using cell2location^56^ (v0.1.2) with annotated scRNA-seq or snRNA-seq references. All datasets are publicly available. For consistency, we removed mitochondrial genes from all spatial datasets prior to deconvolution, as their expression levels are known to be technical artifacts unrelated to cell type abundance. For all scRNA-seq and snRNA-seq datasets, we applied default quality control (QC) thresholds (cell_count_cutoff=5, cell_percentage_cutoff2=0.03, and nonz_mean_cutoff=1.12). Only genes common to both the scRNA-seq/snRNA-seq and spatial datasets were used for deconvolution.

- Mouse hippocampus (scRNA-seq + Slide-seq). We used a preprocessed scRNA-seq dataset from DropViz^57^, and high-resolution Slide-seq data (Puck_200115_08) from Single Cell Portal^14^. Slide-seq data were converted into pseudo-Visium format for validation. Following deconvolution, we performed Leiden clustering on cell type proportion values to define spatial regions for context-specific analysis. Consistent with our previous Slide-seq analyses^27^, both Slide-seq and pseudo-Visium data were restricted to the combined set of the top 2,000 highly variable genes and cell type marker genes prior to CCI analysis. CCI genes identified in the pseudo-Visium from CellNeighborEX v2 were compared to cell contact-dependent genes obtained from the Slide-seq analysis.
- Mouse liver cancer (snRNA-seq + Slide-seq). We utilized paired Slide-seq (mouse_liver_met_2_rna_201002_04) and preprocessed snRNA-seq data from mouse liver cancer^15^. Pseudo-Visium data were generated from Slide-seq. Leiden clustering was performed on deconvolved cell type proportions to define spatial regions used as biological contexts, as spots located in close proximity typically exhibit highly similar cell type compositions. For consistency with the Slide-seq analyses^27^, CCI detection in both datasets was limited to the combined set of the top 2,000 highly variable genes and cell type markers. Validation was performed by assessing the overlap between CCI genes from CellNeighborEX v2 and cell-contact-dependent genes from our previous study.
- Human ovarian cancer (scRNA-seq + Visium + CosMx). We employed paired scRNA-seq (sample Y2, Y3, Y5, MJ10, and MJ11), Visium (sample good_response_P5), and CosMx data from the human ovary (GSE211956)^35^. For scRNA-seq, we followed the cell type annotations provided in the original publication, and for the Visium data, we adopted the malignant cluster labels also provided by the original study^35^. The CosMx data and accompanying scripts were downloaded from the Zenodo repository. We preprocessed the dataset and annotated cell types based on the original scripts. For a fair comparison with CosMx, CCI analysis on the Visium data was restricted to the CosMx gene panel (approximately 980 genes). CosMx data served as a high-resolution benchmark to validate CCI gene predictions derived from the Visium data.
- Human colorectal cancer (scRNA-seq + Visium + Visium HD). We utilized paired scRNA-seq (Chromium Single Cell Flex, aggregated), Visium (Visium CytAssist v2, Sample P2 CRC), and Visium HD with 8 µm spatial resolution (Sample P2 CRC) from human colon^12^. All data were downloaded from the 10x Genomics website. For scRNA-seq, cell type annotations were provided as meta data in the Github repository of original publication^12^. For Visium HD, cell type deconvolution was performed using RCTD^58^ (v2.0.7) based on scripts from the GitHub. We applied Leiden clustering to cell type proportions estimated from Visium data to obtain spatial regions as biological contexts. CCI analysis for both Visium and Visium HD was conducted using all available genes in each dataset (approximately 18,000 genes). From the Visium HD data, we extracted singlets composed of a single cell type and applied CellChat^21^ (v2.1.2) to identify CCI genes. These were then compared to the CCI genes identified by CellNeighborEX v2 in the Visium data.
- Mouse lymph node (scRNA-seq + Visium). We used preprocessed scRNA-seq and Visium data from mouse lymph node^49^. The datasets were obtained from the GitHub repository. Regarding the scRNA-seq, we followed the cell type annotations provided in the original datasets. For both scRNA-seq differential expression analysis and Visium-based CCI gene detection, all genes available in the processed datasets (approximately 11,000 genes) were used without additional filtering.

### RGB-based visualization of interacting cell type pairs

To visually represent spatial patterns of cell-cell interactions, we implemented an RGB-based plotting strategy that captures the relative co-abundance of three interacting cell type pairs within a defined spatial region. Using pseudo-Visium mouse hippocampus data, we selected a cluster enriched for interneurons and identified three biologically relevant neighboring cell types: Cajal-Retzius cells, Entorhinal cells, and Ependymal cells.

For each spot within the target cluster, the interaction strength between interneurons and each neighboring cell type was computed as the product of their respective cell type abundances. These interaction scores were independently normalized to the [0, 1] range to allow comparability across pairs. Each normalized interaction value was then assigned to one of the three color channels: red for Interneuron + Cajal-Retzius, green for Interneuron + Entorhinal, and blue for Interneuron + Ependymal. RGB color values were normalized to ensure that the total intensity per spot remained constrained, thereby enhancing interpretability. This approach enabled intuitive visualization of the spatial co-abundance patterns of distinct interacting cell type pairs within a single spot.

### Validation of spatial proximity-dependent expression

To validate spatial proximity-dependent expression patterns of CCI genes, we used single-molecule spatial transcriptomics data generated by the CosMx platform from a human ovarian cancer tissue sample^35^. We focused on a single field of view (FOV 22), which was annotated with cell type labels and aligned to spatial coordinates.

For each predicted interaction pair consisting of a source cell type and a neighboring cell type, we computed the spatial distances between all source cells and their nearest neighboring cell using the python module of *scipy*.*spatial*.*cKDTree* for a k-d tree algorithm. The cells of the source cell type were divided into “near” and “far” groups based on whether their minimum distance to a neighboring cell was smaller or greater than the median of all such distances. This procedure enabled a consistent and data-driven definition of spatial proximity across all cell type pairs.

We then compared gene expression levels of the predicted CCI gene between the near and far source cell groups. To assess whether expression was significantly increased in spatially proximal cells, we performed one-sided Wilcoxon rank-sum tests (Mann-Whitney U tests), testing the alternative hypothesis that gene expression was higher in near than in far cells. Statistical significance was determined using a threshold of p-value (*p-value* < 0.05). Significant results were visualized with spatial scatter plots and violin plots to illustrate expression differences between proximity-defined groups.

### Validation of CCI genes using conditioned medium experiments

To validate CellNeighborEX v2-specific predictions, we tested whether the expression of selected CCI genes is regulated by interaction-dependent signaling between macrophages and fibroblasts. Specifically, we focused on *CD7* and *SLC11A1*, which were uniquely predicted by CellNeighborEX v2 to be upregulated in macrophages interacting with fibroblasts. Conditioned medium (CM) and Transwell co-culture assays were performed to assess gene expression changes in macrophages in response to fibroblast-derived signals. Detailed protocols for each assay are described below.

#### (i) Cell culture

WI38 and THP-1 cells were cultured in RPMI 1640 media supplemented with 10% FBS and Zell Shield at 37°C in a humidified incubator of 5% CO2. Cells were grown at 2 x 105 cells/mL in 6 well plates. To generate THP-1 derived macrophages, cells were treated with 162nM PMA for 48hr (differentiation) and replaced with fresh media for 48hr (resting).

#### (ii) Co-culture experiments

- Conditioned media (CM) exchange models.

WI38 cells and THP-1 derived macrophages were cultured separately for 48hr. Then, CM was collected from each cell type and subsequently transferred to non-originating cell populations. After incubating with the exchanged CM for 48hr, the cells were harvested.

- Transwell models.

Transwell co-culture system (SPLInsert™ Hanging, 0.4μm, PET, 6-well) was used for this model. After allowing THP-1 cells to rest in fresh media for 48hr, WI38 cells were seeded in the inserts. The co-culture system was incubated for an additional 48hr and the cells were then collected.

#### (iii) Quantitative RT-PCR

Total RNAs were extracted using Trizol (Invitrogen) and reverse-transcribed to cDNA using SuPrimeScript cDNA Synthesis Kit (GeNet Bio). Quantitative RT-PCR was carried out on a BioRad CFX384 with SYBR TOPreal qPCR 2× PreMix (Enzynomics). The relative amounts of mRNA were normalized to β-ACTIN and the fold change of gene expression was calculated using the ddCt method. All samples were run using the following primers: CD7: 5’-AACCTGACTATCACCATGCAC-3’ (F) and 5’-CATCCTTGGGACTGTTCCTC-3’ (R); SLC11A1: 5’-CCATCCCAGACACAAAACCG-3’ (F) and 5’-AGAGAAGTTTGAATCCCGCC-3’ (R); β-ACTIN: 5’-ACCTTCTACAATGAGCTGCG-3’ (F) and 5’-CCTGGATAGCAACGTACATGG-3’ (R).

### Expression correlation analysis using meta-gene scores

To evaluate the biological relevance of CCI genes inferred by CellNeighborEX v2, we performed correlation and enrichment analyses against hallmark pathway gene sets in human colorectal cancer tissue. We focused on two key pathways implicated in tumor-microenvironment (TME) crosstalk: TNF-*α* signaling via NF-κB and KRAS signaling. Corresponding gene sets were downloaded from the MSigDB v2024.1 mouse hallmark collection.

Gene expression matrices from Visium data were first normalized to a total count of 10,000 per spot and log-transformed using scanpy.pp.normalize_total and scanpy.pp.log1p. To summarize the expression of each gene set across spatial spots, we computed meta-gene scores using the *score_genes()* function in Scanpy. For a given gene set *G*, the meta-gene score for spot *s* is defined as:

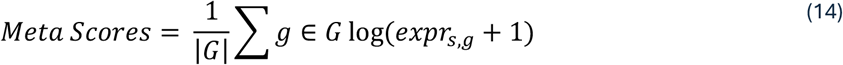

where *expr*_*s,g*_ represents the normalized expression level of gene *g* at spot *s*, and |*G*| denotes the number of genes in the set. No control gene set was subtracted in our scoring (*i*.*e*., ctrl_size=None).

For each spatial region, CCI gene sets identified by CellNeighborEX v2 were used to compute additional meta-gene scores. To evaluate the association between CCI gene expression and pathway activity, we computed Spearman’s rank correlation coefficient between the CCI score and each pathway score across all spots within the same cluster. In parallel, Fisher’s exact test was used to assess the enrichment of TNF-*α* and KRAS pathway genes within each cluster-specific CCI gene set, using the set of all expressed genes as the background.

Statistical significance of correlations and enrichments was adjusted using Bonferroni correction. The results were visualized using bar plots of Spearman’s ρ with significance annotations and - log^10^-transformed p-values for Fisher’s exact test.

### Survival analysis of *FOXP3* expression

To investigate the clinical relevance of *FOXP3* expression, we performed survival analysis using RNA-seq and clinical data from The Cancer Genome Atlas (TCGA) colorectal cancer cohort, including both colon (COAD) and rectal (READ) adenocarcinoma patients. Gene expression quantification data (STAR - Counts) and corresponding clinical data (patient XML files) were retrieved using the TCGAbiolinks package (v2.26.0). *FOXP3* expression values were extracted from the processed count matrix, and samples were stratified into “High” and “Low” expression groups based on the median expression level of *FOXP3* across all patients.

Overall survival (OS) was defined as the number of days from diagnosis to death or last followup. For patients who were alive at the last follow-up, OS time was defined using days_to_last_followup. The primary endpoint was overall survival status (vital_status), coded as 1 (deceased) or 0 (censored). Expression and survival data were merged by patient barcode (first 12 characters of TCGA sample ID).

Kaplan-Meier survival curves were generated using the survival and survminer packages in R. Differences between survival curves were assessed using the log-rank test. The survival object was defined as Surv (OS_time, OS_event), and group-wise survival curves were estimated using the survfit() function. Visualization was performed with ggsurvplot(), displaying survival probability over time for *FOXP3*-high and *FOXP3*-low groups. The log-rank test p-value was reported to assess the statistical significance of survival differences between the two groups.

### Functional validation and enrichment analyses

To evaluate the functional relevance of experimental condition-specific CCI genes identified by CellNeighborEX v2, we performed enrichment and semantic similarity analyses using Visium data from murine lymph nodes under Mycobacteria-infected (MS) and control (PBS) conditions. These analyses aimed to determine whether MS-specific CCI genes reflect biologically meaningful immune responses, such as activation of the interferon gamma (IFN-γ) pathway.

We first constructed a ranked gene list based on log^2^ fold changes (logFC) between MS and PBS samples, calculated from the log-normalized Visium expression matrix. For each gene *g*, logFC was computed as:

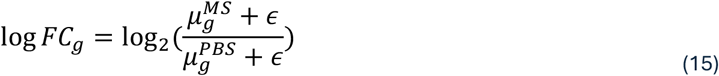

where 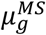 and 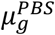 are average expression values of gene *g* , across MS and PBS spots, respectively, and *ϵ* = 1 × 10^−5^ is a pseudocount to avoid division by zero. The resulting pre-ranked gene list served as input to Gene Set Enrichment Analysis (GSEA) using the fgsea R package with 10,000 permutations.

We obtained IFN-γ response gene set from the MSigDB (v2024.1, mouse version). GSEA was used to assess the enrichment of IFN-γ response genes in the MS condition. We then mapped the ranks of MS-specific and PBS-specific CCI genes onto the GSEA-ranked gene list and visualized their distribution.

To quantitatively examine the functional similarity between CCI genes and the IFN-γ response program, we computed semantic similarity scores using Gene Ontology (GO) Biological Process annotations. CCI genes and IFN-γ response genes were converted to Entrez Gene IDs, and only genes with GO annotations were retained. We then calculated pairwise GO term-based semantic similarity using the Wang method^59^, as implemented in the GOSemSim R package^60^. For each CCI gene, a similarity score was computed as the average similarity to all IFN-γ response genes using the Best Match Average (BMA) strategy^61^.

## Data availability

- Slide-seq V2 data in a mouse hippocampus: Single Cell Portal (https://singlecell.broadinstitute.org/single_cell/study/SCP815/sensitive-spatial-genome-wide-expression-profiling-at-cellular-resolution#study-summary).
- scRNA-seq data in mouse hippocampus: DropViz (http://dropviz.org/).
- Slide-seq and snRNA-seq data in mouse liver cancer: Single Cell Portal (https://singlecell.broadinstitute.org/single_cell/study/SCP1278/spatial-genomics-enables-multi-modal-study-of-clonal-heterogeneity-in-tissues).
- scRNA-seq and Visium data in human ovarian cancer: GSE211956 (https://www.ncbi.nlm.nih.gov/geo/query/acc.cgi?acc=GSE211956)
- CosMx data in human ovarian cancer: Zenodo (https://zenodo.org/records/10048057)
- scRNA-seq, VisiumHD, and Visium data in human colon cancer: 10X Genomics (https://www.10xgenomics.com/platforms/visium/product-family/dataset-human-crc)
- scRNA-seq and Visium data in mouse lymph node: Github (https://github.com/romain-lopez/DestVI-reproducibility/tree/master/lymph_node/deconvolution)
- Precomputed CellNeighborEX v2 outputs (sc_ccisignal.h5ad, sp_ccisignal.h5ad): figshare (https://figshare.com/s/a22ae3508f9ea0b091c3).

## Code availability

- The source code for CellNeighborEX v2 is available at: GitHub (https://github.com/hkim240/CellNeighborEX-v2)
- CosMx preprocessing and cell type annotations for human ovarian cancer were performed using publicly available resources:
- Zenodo (https://zenodo.org/records/10048057)
- scRNA-seq annotation and Visium HD deconvolution for human colon cancer were performed using publicly available code:

Github (https://github.com/10XGenomics/HumanColonCancer_VisiumHD)

