## Supplementary Information for "Identifying Context-Specific Cell-Cell Interaction Genes Without Ligand-Receptor Databases from Spatial Transcriptomics"

This PDF file includes Supplementary Figures 1-20.

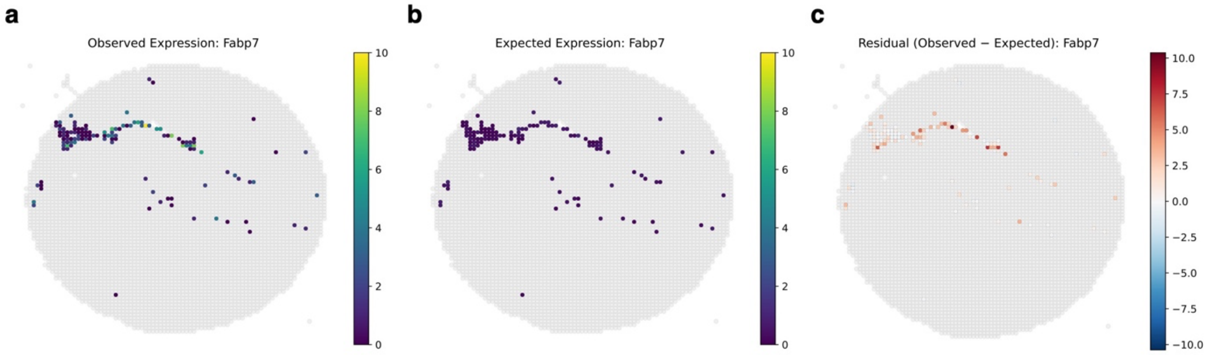


**Supplementary Fig. 1. Comparison of observed, expected, and residual spatial expression patterns for *Fabp7* in pseudo-Visium hippocampus data. a** Observed expression of *Fabp7*, showing localized enrichment in specific spatial regions. **b** Expected expression of *Fabp7* estimated from scRNA-seq reference profiles. The expected signal remains low and relatively uniform across the astrocyte-containing spots, reflecting the dilution of interaction-specific signals in population-level scRNA-seq data. **c** Residual expression (Observed - Expected) of *Fabp7*. By subtracting the reference-derived expectation, CellNeighborEX v2 recovers spatially restricted hotspots where *Fabp7* expression is significantly enhanced beyond the predicted levels. These high-residual regions identify putative sites of astrocyte-endothelial tip cell interaction, consistent with previous co-culture assays demonstrating that *Fabp7* is specifically induced in astrocytes upon contact with endothelial tip cells. Only spots belonging to the relevant cluster (Cluster 13) are displayed, with other spots indicated in light gray.

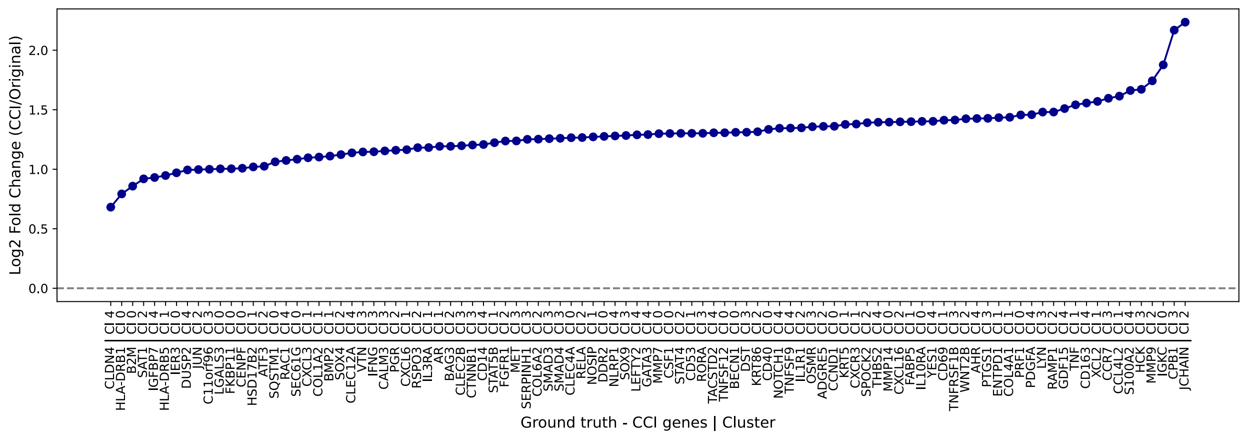


**Supplementary Fig. 2. Simulated log fold changes for ground-truth CCI genes in synthetic benchmark.** To mimic biologically plausible CCI-driven expression, synthetic expression signals were added to 99 ground-truth CCI genes based on the co-abundance of paired source and neighboring cell types within spatial spots. The resulting log fold changes ranged from 0.5 to 2.5, capturing realistic levels of transcriptional variation. Of the 99 genes, 69 were annotated in existing interaction databases (Omnipath, CellChat, CellTalk), while 30 were not, providing a balanced and challenging evaluation framework.

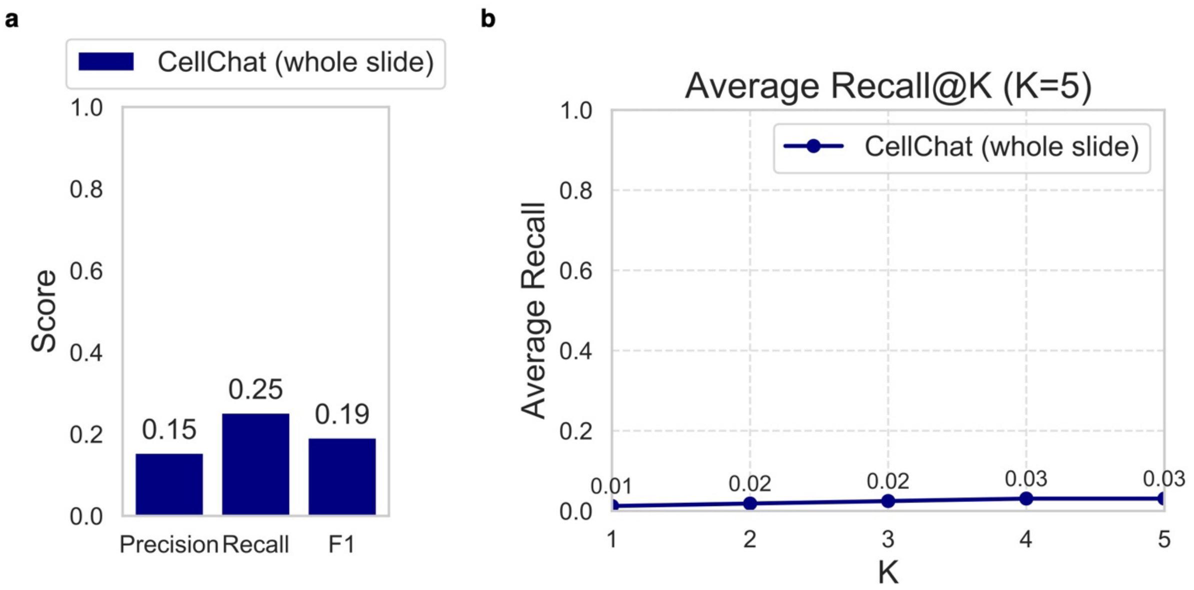


**Supplementary Fig. 3. Benchmarking CellChat on synthetic Visium data. a** Gene-level performance of CellChat on the synthetic dataset, showing precision, recall, and F1 scores. CellChat exhibited low values across all three metrics, reflecting poor recovery of true CCI genes. **b** Accuracy of interacting cell type pair attribution by CellChat, measured by Recall@5. The low Recall@5 values indicate limited ability to identify correct source-neighboring cell type relationships.

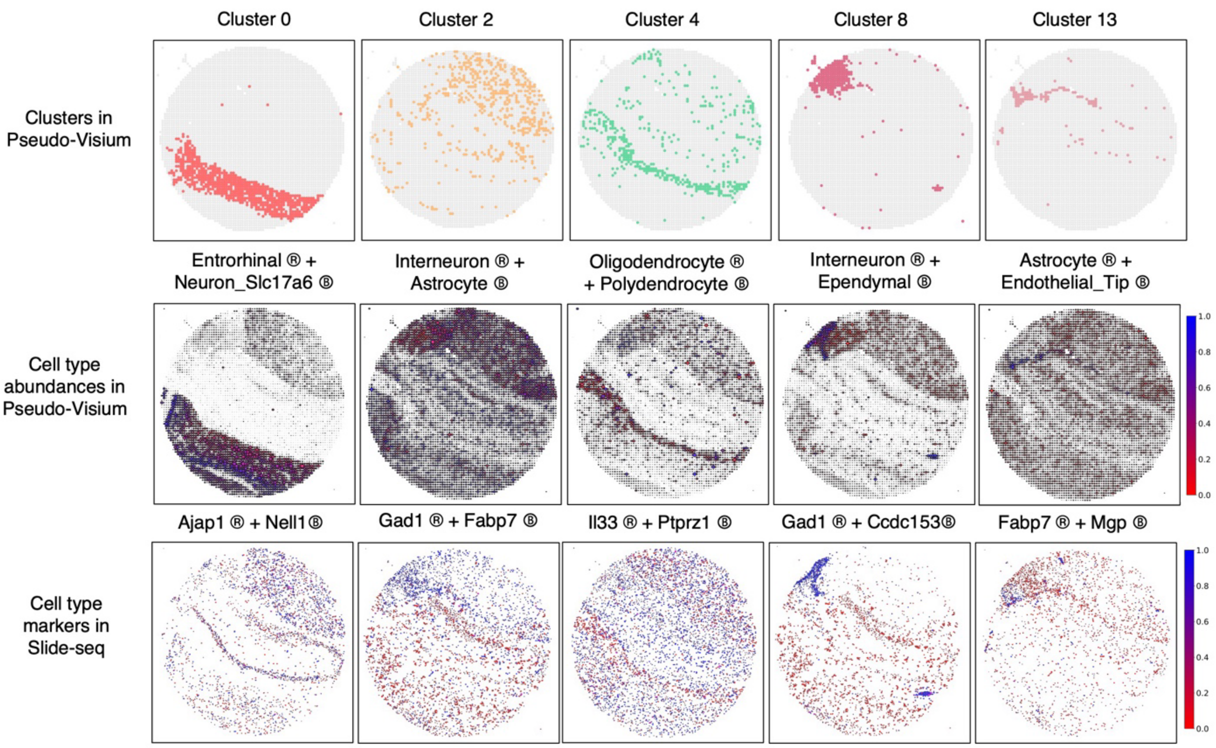


**Supplementary Fig. 4. Spatial colocalization of predicted interacting cell type pairs in mouse hippocampus Slide-seq data.** For each spatial cluster identified in the pseudo-Visium dataset, CellNeighborEX v2 inferred CCI genes and corresponding source-neighbor cell type pairs. To validate these predictions, we examined the original high-resolution Slide-seq data in mouse hippocampus and confirmed the co-presence of the predicted cell types. Spatial colocalization was supported by the expression of canonical marker genes specific to the inferred cell types: *Ajap1* (Entorhinal), *Nell1* (Neuron_Slc17a6), *Gad1* (Interneuron), *Fabp7* (Astrocyte), *Il33* (Oligodendrocyte), *Ptprz1* (Polydendrocyte), *Ccdc153* (Ependymal), and *Mgp* (Endothelial_Tip). Representative marker genes for each cell type pair showed spatial co-expression, supporting the plausibility of the inferred interactions.

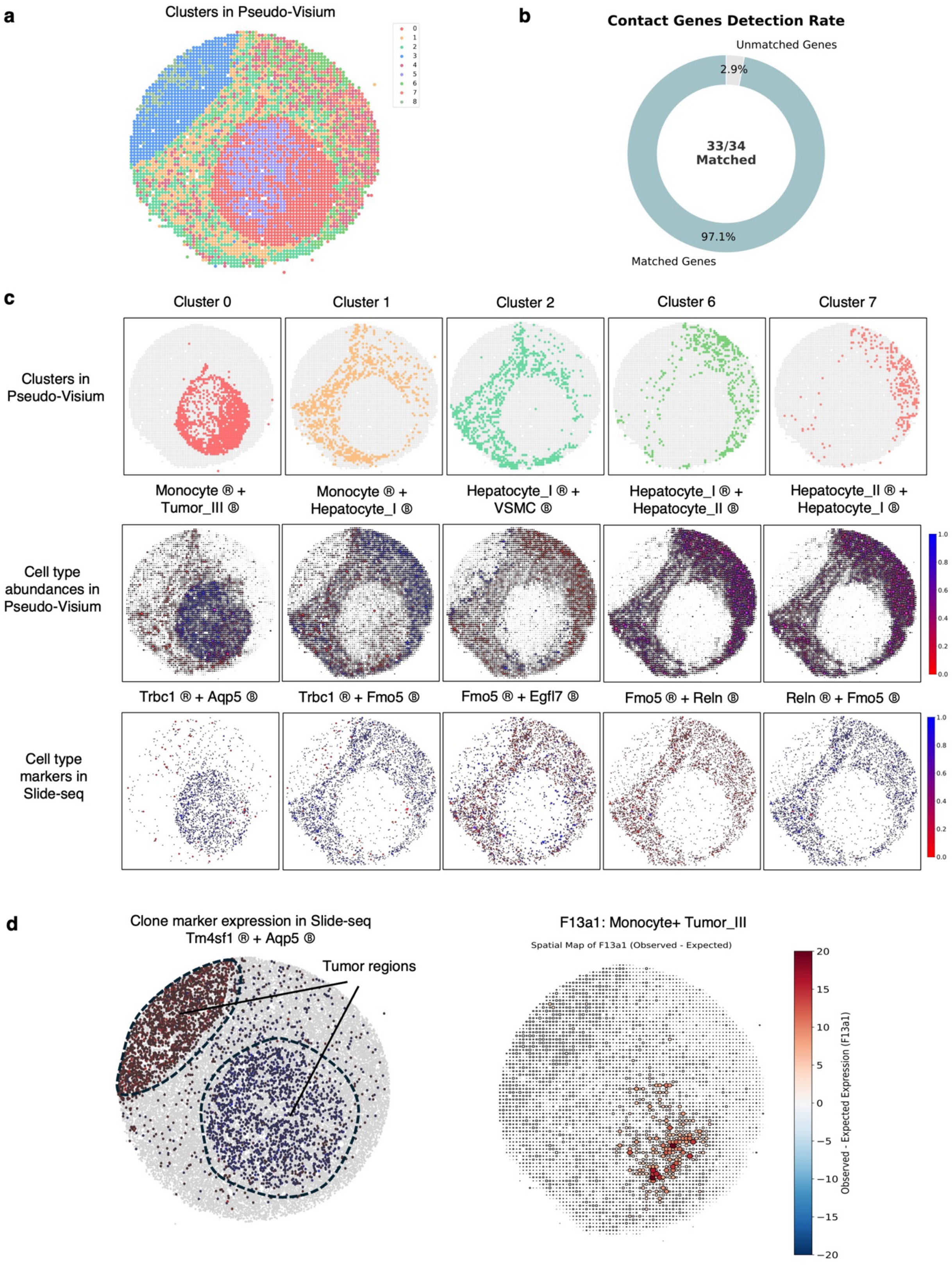

a
Clusters in Pseudo-Visium
b
Contact Genes Detection Rate
Unmatched Genes
2.9%
33/34
Matched
97.1%
Matched Genes
c
Cluster 0
Cluster 1
Cluster 2
Cluster 6
Cluster 7
Clusters in Pseudo-Visium
Monocyte ® + Tumor_III Ⓑ
Monocyte ® + Hepatocyte_I Ⓑ
Hepatocyte_I ® + VSMC Ⓑ
Hepatocyte_I ® + Hepatocyte_II Ⓑ
Hepatocyte_II ® + Hepatocyte_I Ⓑ
Cell type abundances in Pseudo-Visium
Trbc1 ® + Aqp5 Ⓑ
Trbc1 ® + Fmo5 Ⓑ
Fmo5 ® + Egfl7 Ⓑ
Fmo5 ® + Reln Ⓑ
Reln ® + Fmo5 Ⓑ
Cell type markers in Slide-seq
d
Clone marker expression in Slide-seq
Tm4sf1 ® + Aqp5 Ⓑ
Tumor regions
F13a1: Monocyte+ Tumor_III
Spatial Map of F13a1 (Observed - Expected)
Observed - Expected Expression (F13a1)

**Supplementary Fig. 5. Validation of CellNeighborEX v2 predictions in mouse liver cancer using Slide-seq. a** Spatial clustering of pseudo-Visium data aggregated from Slide-seq reveals nine distinct spatial regions. **b** CellNeighborEX v2 successfully recovered 33 of 34 previously identified contact-mediated genes (97.1% match). **c** Representative spatial clusters from pseudo-Visium (top), predicted source-neighbor cell type pairs and their co-abundance (middle), and spatial expression of corresponding marker genes in Slide-seq (bottom). *Trbc1* (monocyte marker) and *Aqp5* (tumor cell subtype III marker) are co-expressed in regions where monocytes and tumor cells interact. Other examples include *Fmo5* (hepatocyte I marker), *Egfl7* (vascular smooth muscle cell, VSMC marker), and *Reln* (hepatocyte II marker), which show spatial co-expression with their predicted partners, supporting interaction predictions. **d** Spatial definition of tumor regions and localization of interaction-associated *F13a1* residual signals. Tumor-specific regions were defined using matched Slide-seq data based on the spatial expression of *Tnfaip1* and *Aqp5*, known markers for tumor cells (left). CCI expression of *F13a1* (i.e., observed - expected expression) in pseudo-Visium data is enriched in monocytes interacting with tumor cells (right). Spot size is scaled proportionally to the absolute magnitude of the residual.

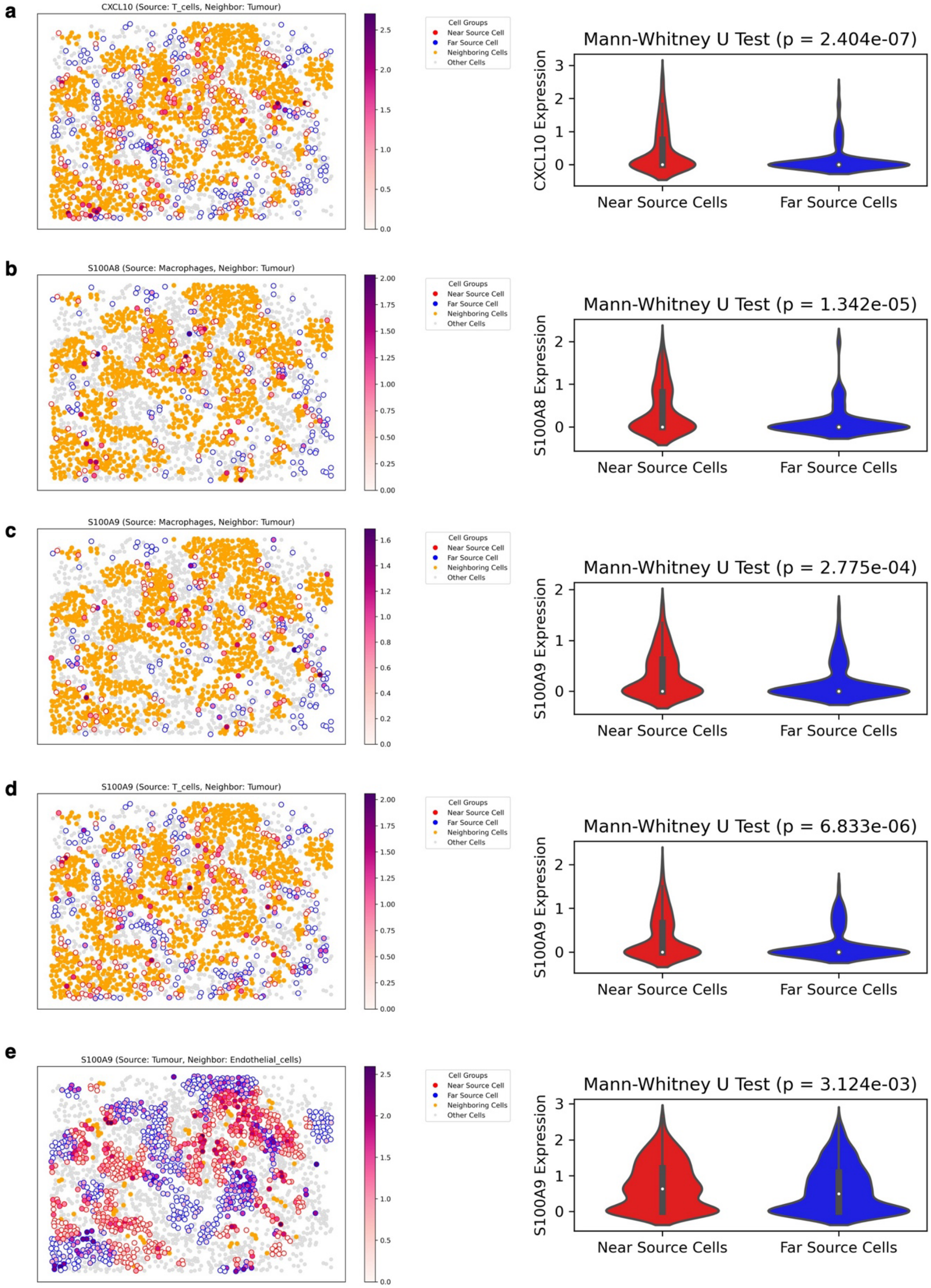
a
CXCL10 (Source: T_cells, Neighbor: Tumour)
Cell Groups
Near Source Cell
Far Source Cell
Neighboring Cells
Other Cells
Mann-Whitney U Test (p = 2.404e-07)
CXCL10 Expression
Near Source Cells
Far Source Cells
b
S100A8 (Source: Macrophages, Neighbor: Tumour)
Cell Groups
Near Source Cell
Far Source Cell
Neighboring Cells
Other Cells
Mann-Whitney U Test (p = 1.342e-05)
S100A8 Expression
Near Source Cells
Far Source Cells
c
S100A9 (Source: Macrophages, Neighbor: Tumour)
Cell Groups
Near Source Cell
Far Source Cell
Neighboring Cells
Other Cells
Mann-Whitney U Test (p = 2.775e-04)
S100A9 Expression
Near Source Cells
Far Source Cells
d
S100A9 (Source: T_cells, Neighbor: Tumour)
Cell Groups
Near Source Cell
Far Source Cell
Neighboring Cells
Other Cells
Mann-Whitney U Test (p = 6.833e-06)
S100A9 Expression
Near Source Cells
Far Source Cells
e
S100A9 (Source: Tumour, Neighbor: Endothelial_cells)
Cell Groups
Near Source Cell
Far Source Cell
Neighboring Cells
Other Cells
Mann-Whitney U Test (p = 3.124e-03)
S100A9 Expression
Near Source Cells
Far Source Cells

**Supplementary Fig. 6. Spatial proximity-dependent expression of predicted CCI genes in CosMx data in human ovarian cancer.** For each predicted cell-cell interaction, source cells were stratified into groups based on their spatial proximity to the predicted neighboring cell type. Gene expression was compared between source cells that were spatially proximal versus those distant from their interacting partners using Mann-Whitney U tests. **a-e** *CXCL10* (a), *S100A8* (b), and *S100A9* (c–e) exhibit significantly higher expression in source cells when positioned near their predicted interacting cell types. Specifically, *CXCL10* is highly expressed in T cells adjacent to tumor cells (a); *S100A8* and *S100A9* are increased in macrophages near tumor cells (b, c); *S100A9* is also upregulated in T cells near tumor cells (d) and in tumor cells close to endothelial cells (e).

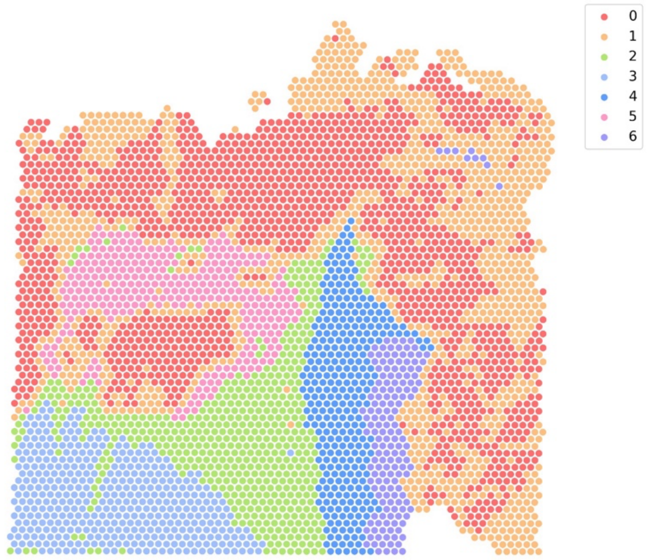


**Supplementary Fig. 7. Spatial clustering of Visium data from human colorectal cancer tissue.** Seven spatial clusters were identified from the Visium dataset. Each hexagonal spot represents a Visium spot, colored according to its assigned spatial cluster. These clusters served as the basis for detecting spatial region-specific cell-cell interaction (CCI) genes using CellNeighborEX v2.

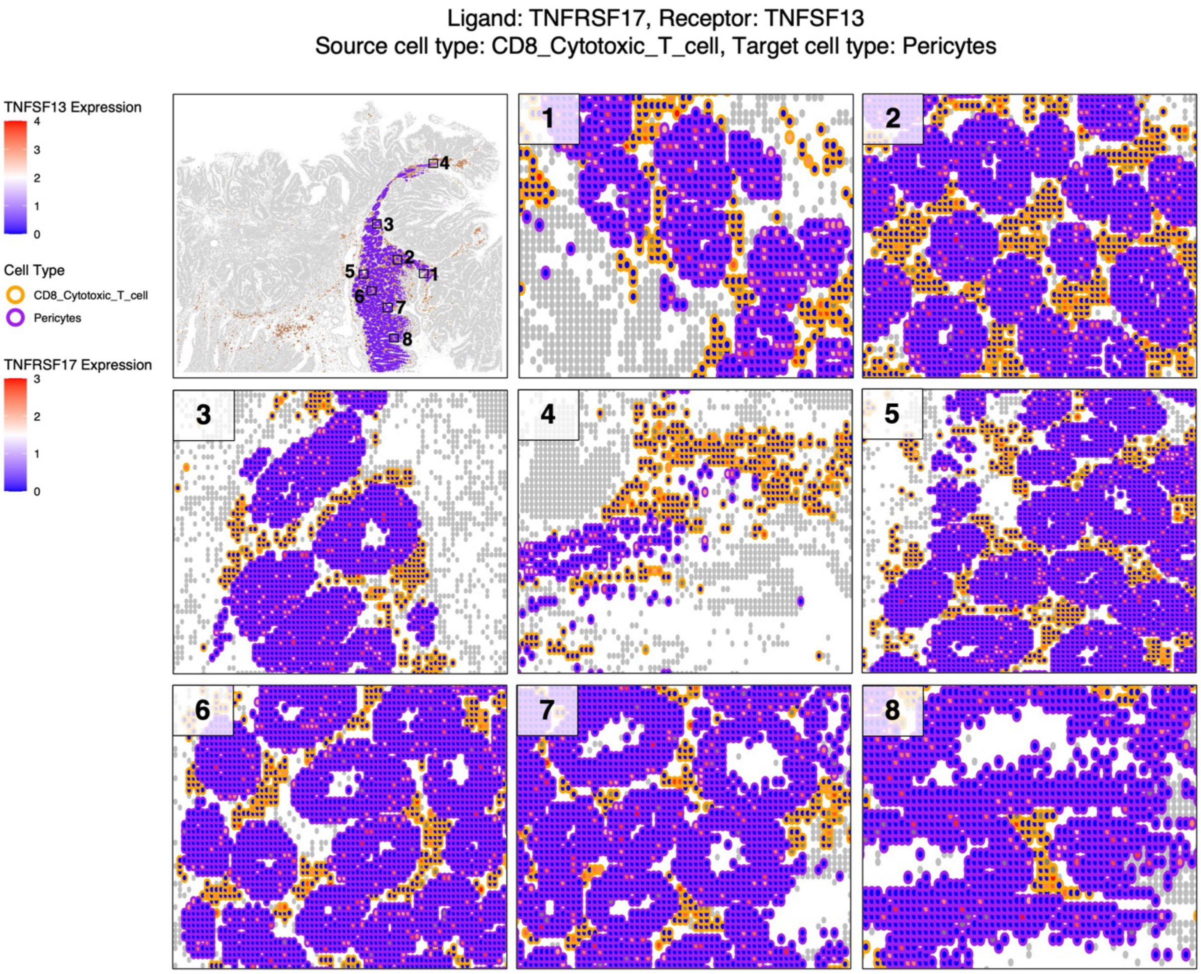


**Supplementary Fig. 8. Spatial validation of the *TNFRSF17-TNFSF13* interaction in human colorectal cancer Visium HD data.** Eight fields of view corresponding to cluster 4 in the Visium data were examined. CD8$^+$ cytotoxic T cells (orange; source cell type) and pericytes (purple; neighboring cell type) were co-localized across these regions. The predicted ligand *TNFRSF17* was highly expressed in the source cell type, while the receptor *TNFSF13* was expressed in the neighboring cell type. These spatial patterns in the high-resolution Visium HD data support the interaction inferred by CellNeighborEX v2 from the low-resolution Visium data.

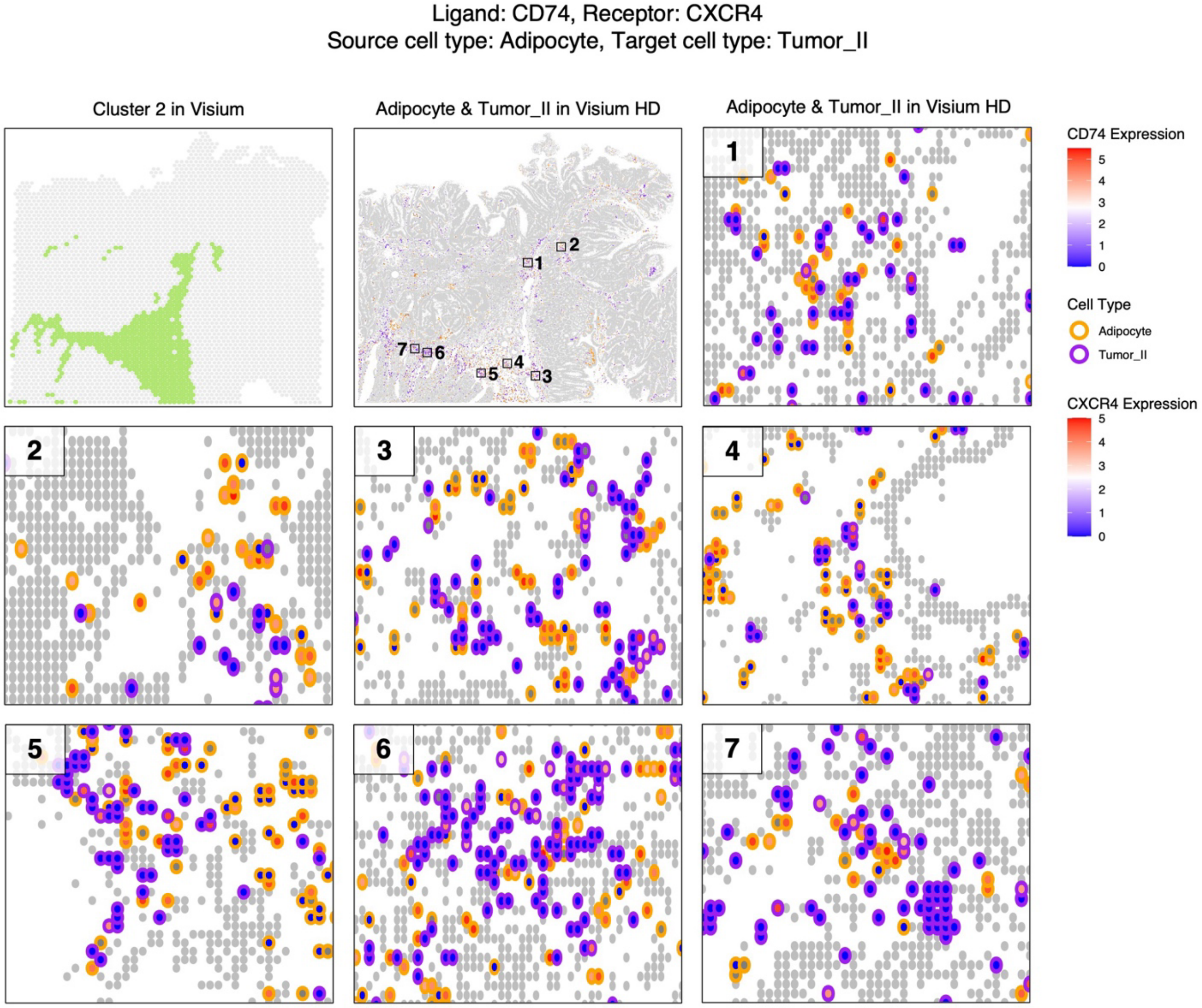


**Supplementary Fig. 9. Spatial validation of the *CD74-CXCR4* interaction in human colorectal cancer Visium HD data.** Seven fields of view corresponding to cluster 2 in the Visium data were examined. Adipocytes (orange; source cell type) and Tumor_II cells (purple; neighboring cell type) were co-localized in these regions. The predicted ligand *CD74* was highly expressed in the source cell type, and the receptor *CXCR4* was expressed in the neighboring cell type. These spatial co-expression patterns provide supporting evidence for the interaction predicted by CellNeighborEX v2.

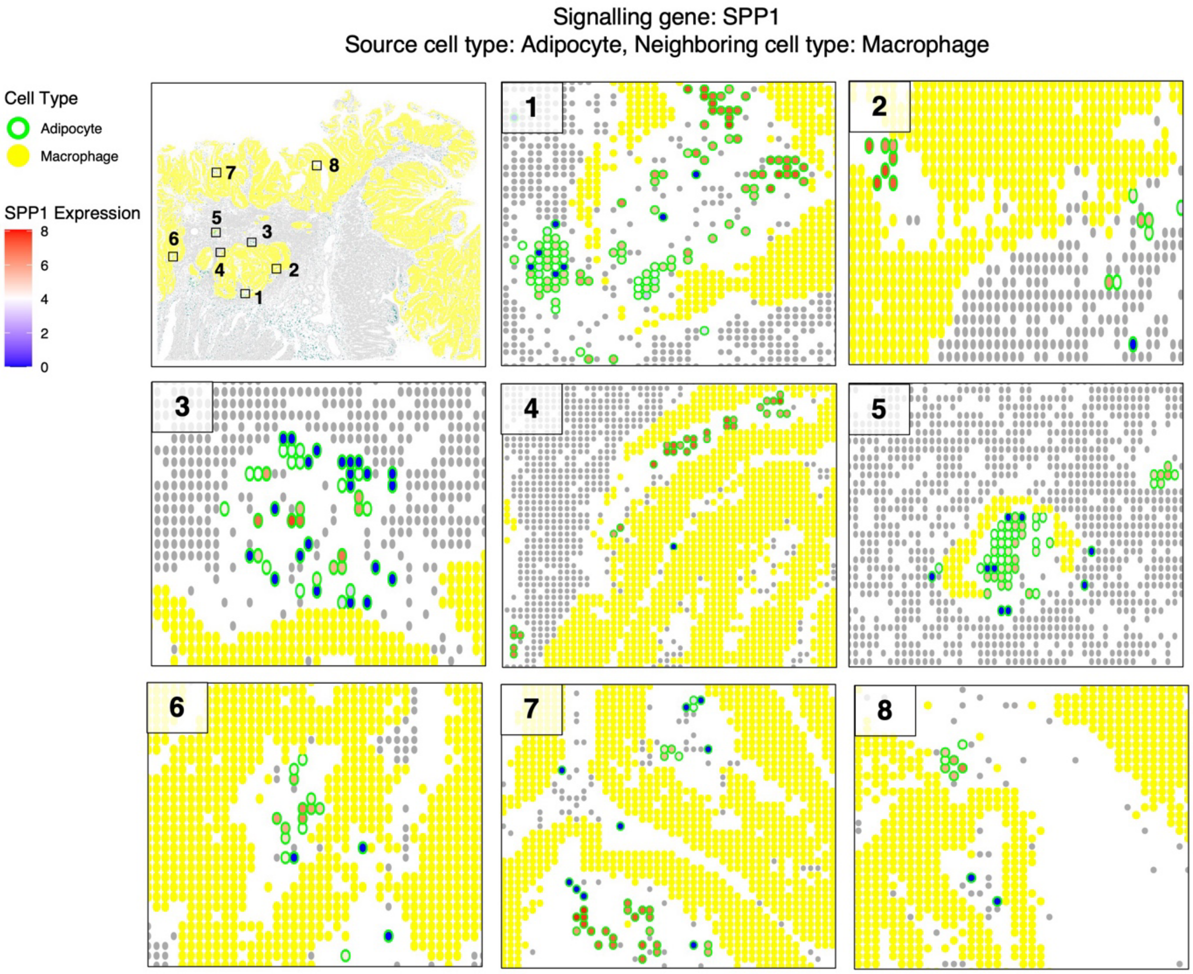


**Supplementary Fig. 10. Recovery of CellChat-predicted signaling gene *SPP1* by CellNeighborEX v2.** CellChat identified *SPP1* as a signaling gene mediating interactions from adipocytes (source cell type) to macrophages (neighboring cell type) in the Visium HD data. CellNeighborEX v2, when applied to the matched low-resolution Visium dataset, recovered this gene and cell type pair as a spatial region-specific CCI. Spatial mapping in Visium HD confirms the co-localization of adipocytes and macrophages, with strong *SPP1* expression observed in their shared regions.

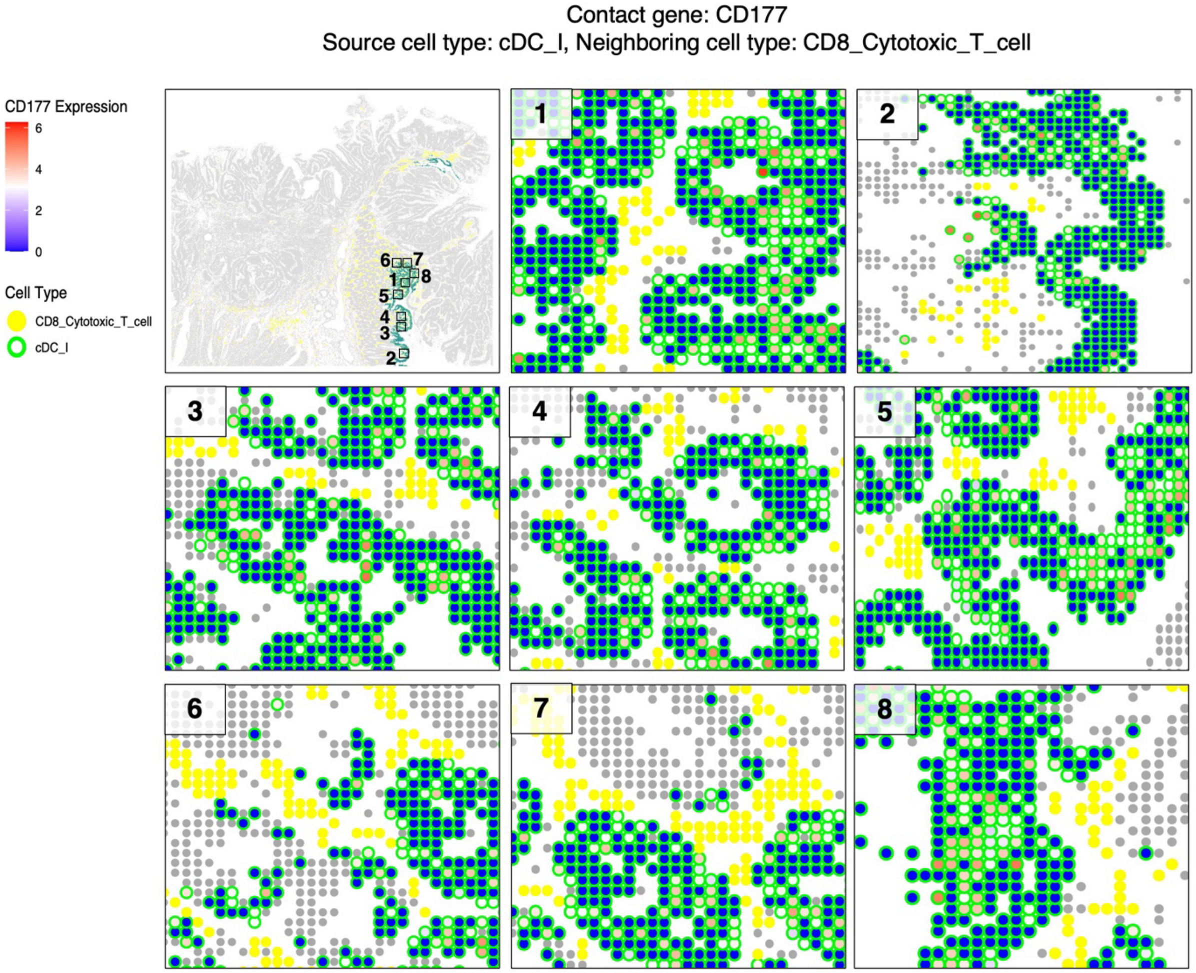


**Supplementary Fig. 11. Recovery of CellChat-predicted contact gene *CD177* by CellNeighborEX v2.** *CD177* was predicted by CellChat as a contact-mediated CCI gene between cDC_I (source cell type) and CD8_Cytotoxic_T_cells (neighboring cell type) in Visium HD data. CellNeighborEX v2 recovered this gene-cell pair in the Visium dataset. Spatial visualization in Visium HD shows *CD177* expression enriched in regions where the predicted interacting cell types co-occur.

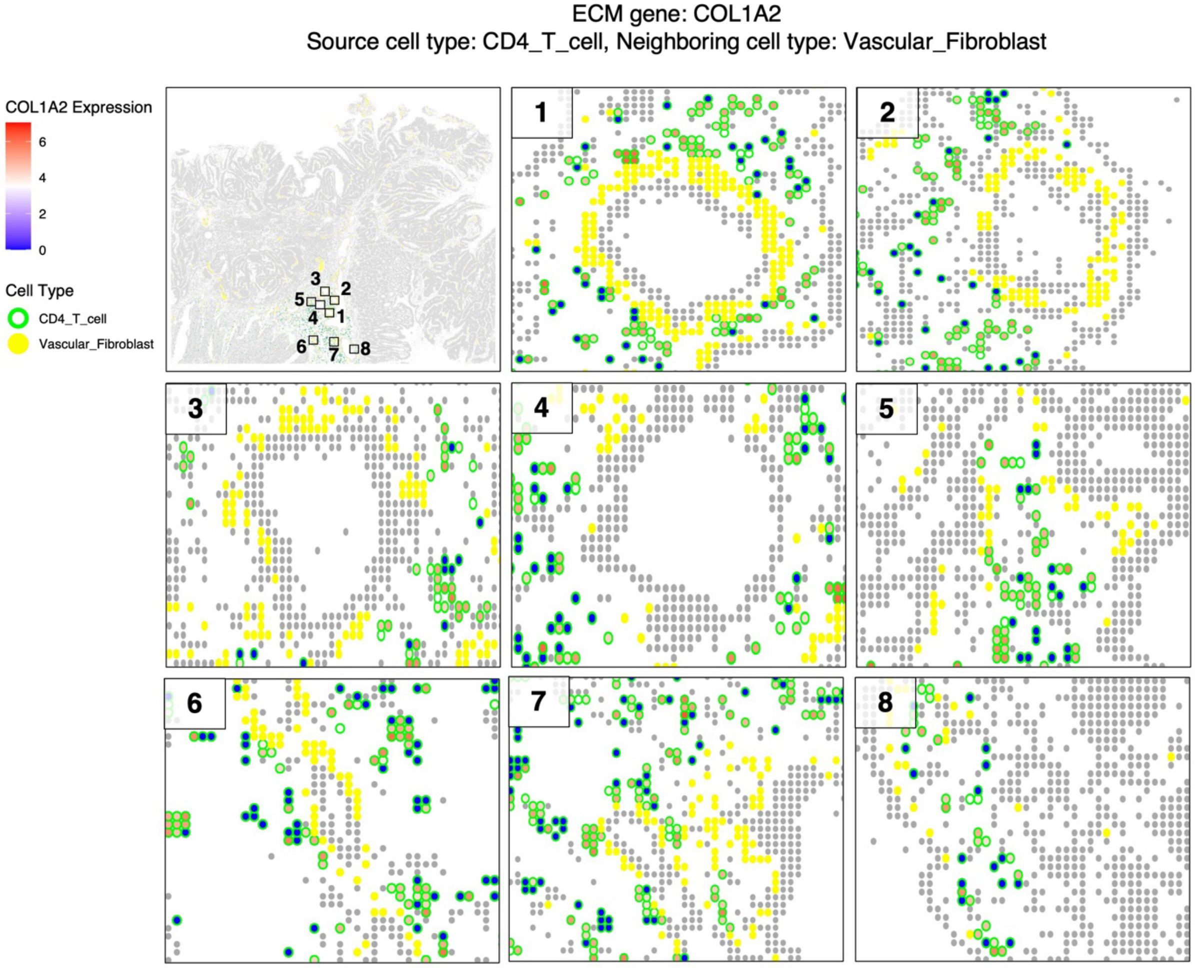


**Supplementary Fig. 12. Recovery of CellChat-predicted ECM gene *COL1A2* by CellNeighborEX v2.** *COL1A2* was identified by CellChat as an ECM-related gene involved in communication from CD4_T_cells (source cell type) to Vascular_Fibroblasts (neighboring cell type) in the Visium HD dataset. This gene and cell pair were recovered by CellNeighborEX v2 when applied to the corresponding Visium data. Spatial maps in Visium HD display co-localization of the two cell types and increased *COL1A2* expression in the interacting regions.

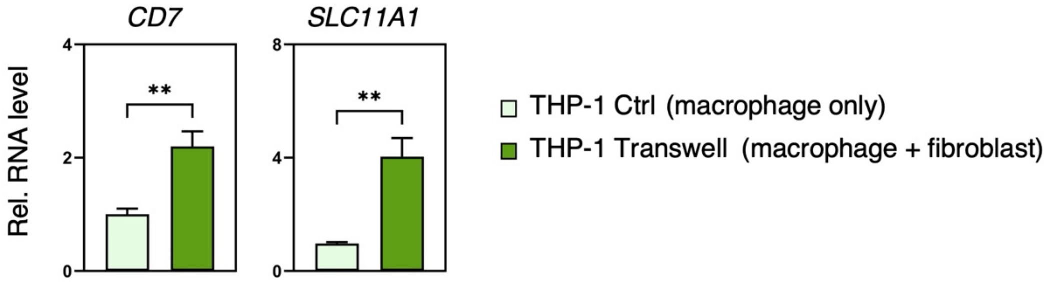


**Supplementary Fig. 13. Transwell validation of interaction-driven expression of *CD7* and *SLC11A1* in macrophages.** To validate CellNeighborEX v2-specific predictions, we performed Transwell co-culture experiments in which THP-1-derived macrophages were cultured with or without WI38 fibroblasts, separated by a permeable membrane. Quantitative RT-PCR analysis revealed that co-culture with fibroblasts significantly increased the expression of *CD7* and *SLC11A1* in macrophages, supporting interaction-dependent induction. *CD7* expression was specific to macrophages. Data are presented as mean ± SE (n = 3). Statistical significance was assessed using a two-tailed Student's t-test.

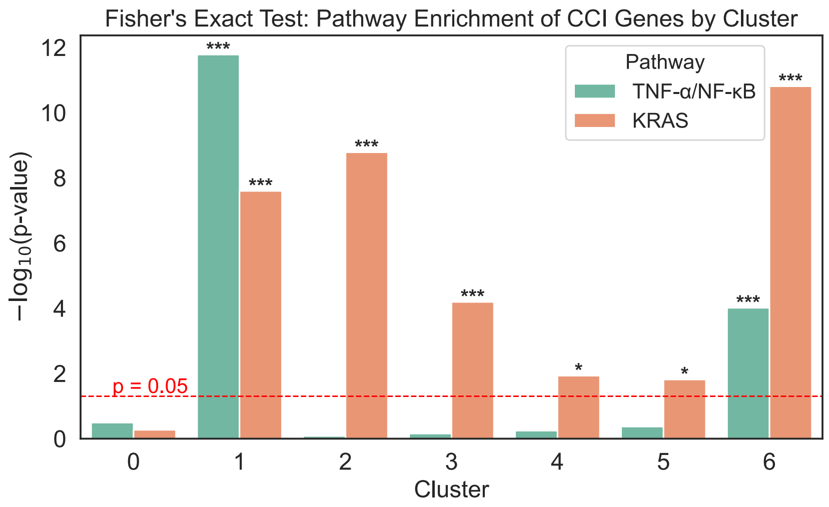


**Supplementary Fig. 14. Pathway enrichment of CellNeighborEX v2-inferred CCI genes across Visium clusters in human colorectal cancer.** Fisher's exact test was used to assess the enrichment of TNF-α/NF-κB (green) and KRAS (orange) signaling pathway genes among CellNeighborEX v2-inferred CCI genes in each of the seven spatial clusters from the Visium dataset. The red dashed line marks the significance threshold ($p = 0.05$). Cluster 1 exhibited the most prominent co-enrichment of both pathways. Asterisks indicate significance levels (*$p < 0.05$, ***$p < 0.001$).

a

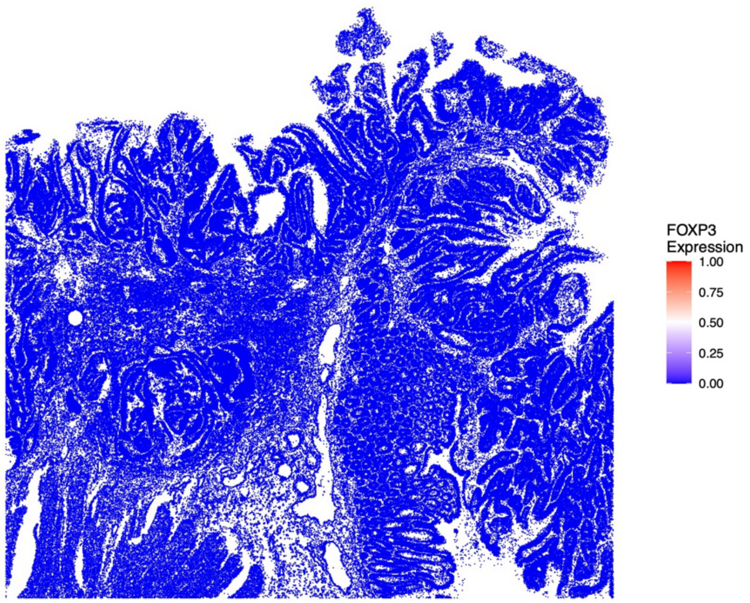


b

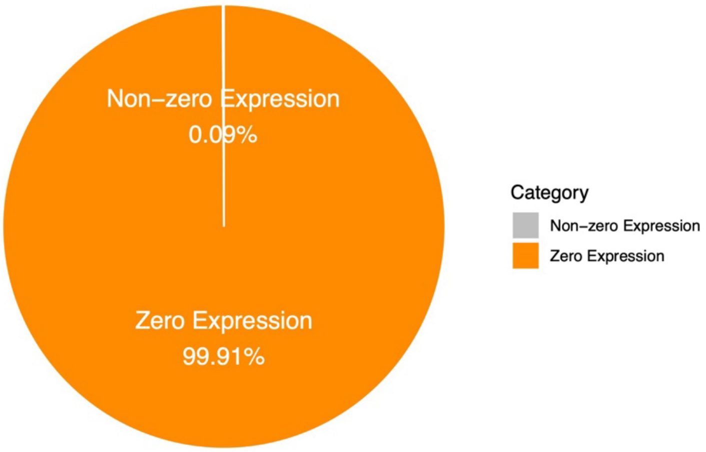


**Supplementary Fig. 15. Limited detection of *FOXP3* in human colorectal cancer Visium HD dataset. a** Spatial expression map of *FOXP3* across Visium HD spots, showing near-complete absence of signal throughout the tissue section. **b** Pie chart quantifying *FOXP3* expression levels across all spots, revealing that 99.91% of spots exhibit zero expression, and only 0.09% show any detectable transcripts. These results illustrate the low sensitivity of high-resolution spatial transcriptomics platforms for detecting low-abundance genes such as *FOXP3*.

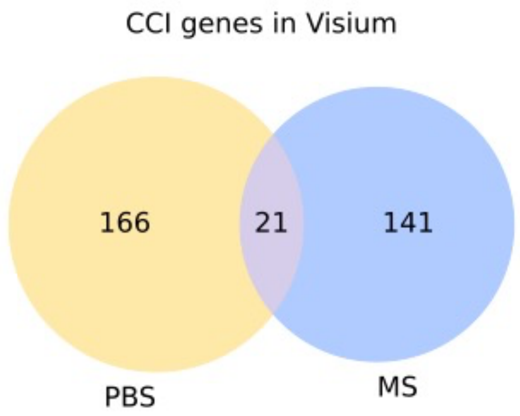


**Supplementary Fig. 16. Condition-specific CCI genes in mouse lymph node Visium data.** Venn diagram showing the overlap of CCI genes inferred by CellNeighborEX v2 between PBS control and Mycobacteria (MS)-infected lymph node samples. A total of 166 and 141 genes were uniquely detected in PBS and MS conditions, respectively, with 21 CCI genes shared between both conditions.

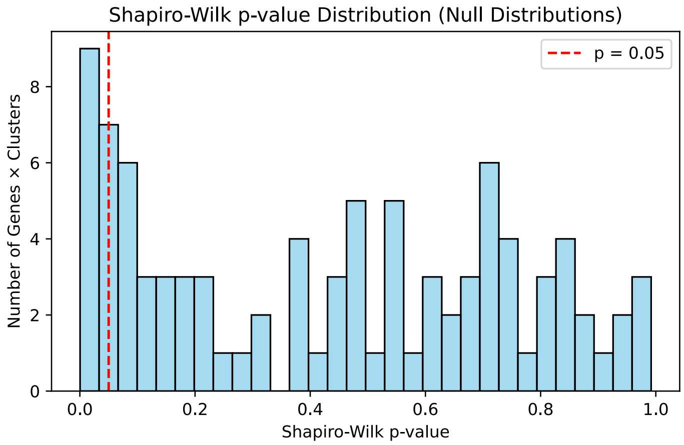


**Supplementary Fig. 17. Normality assessment of null distributions for permutation-based context specificity test.** Histogram showing the distribution of Shapiro-Wilk p-values for permutation-derived null distributions across gene-cluster combinations in the synthetic dataset. The majority of null distributions passed the normality test ($p > 0.05$; red dashed line), indicating that Z-score-based inference is valid for most gene-context pairs. Less than 10% of combinations showed significant deviation from normality.

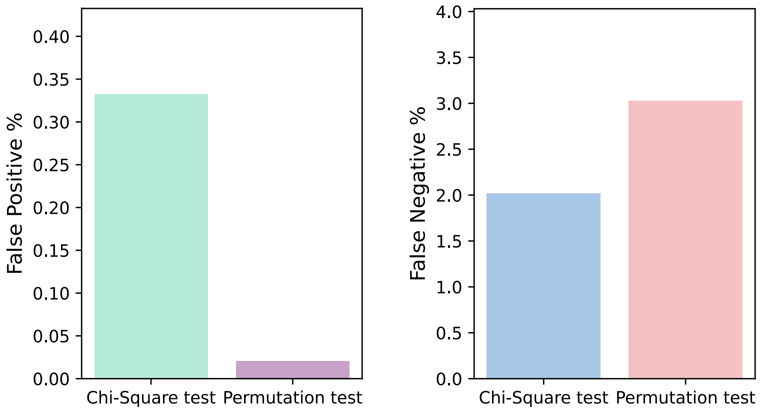


**Supplementary Fig. 18. Performance comparison between chi-squared test and permutation test in the synthetic dataset.** Left: False positive rates for CCI gene detection were substantially lower when using the permutation test compared to the chi-squared test. Right: Conversely, the chi-squared test demonstrated a lower false negative rate than the permutation test. These results highlight the tradeoff between specificity and sensitivity when using each test independently.

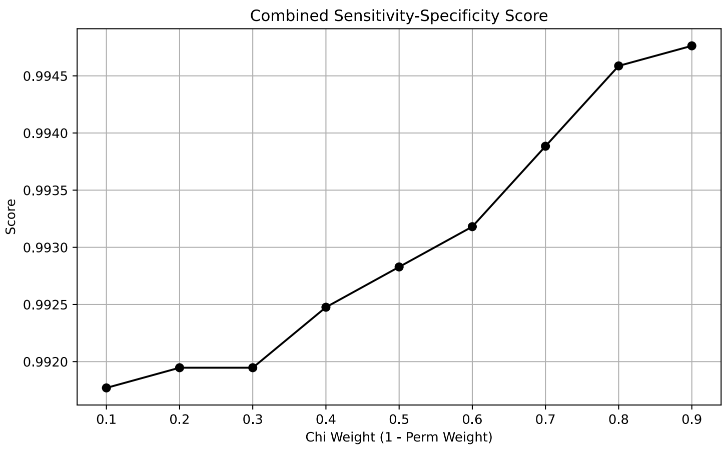


**Supplementary Fig. 19. Evaluation of combined sensitivity-specificity scores across different weighting schemes for Cauchy p-value integration.** The harmonic mean of sensitivity and specificity was computed for each weight combination of chi-squared and permutation p-values. A 0.9 (chi-squared) to 0.1 (permutation) weighting yielded the highest combined score, indicating the optimal tradeoff between detection sensitivity and false positive control.

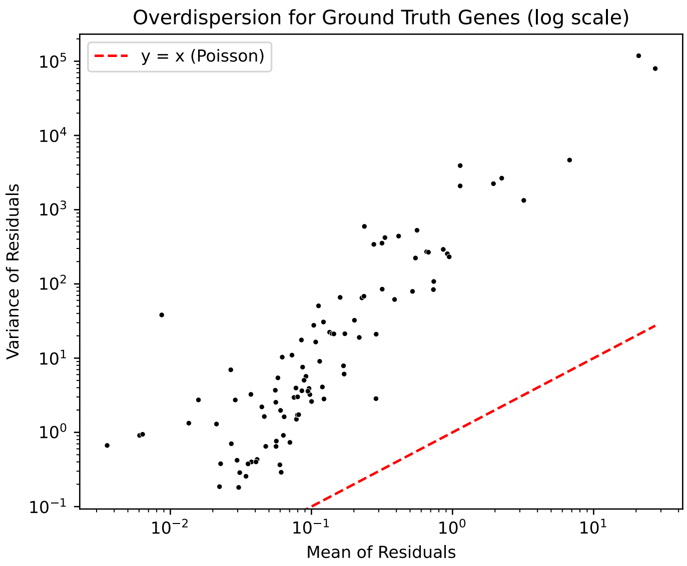


**Supplementary Fig. 20. Overdispersion of residual expression values for ground truth CCI genes in the synthetic dataset.** The scatter plot shows the mean versus variance of residuals across spatial spots for each ground truth gene on a log-log scale. The red dashed line represents the Poisson expectation (variance = mean). Most points lie above the line, indicating substantial overdispersion in residuals, which supports the appropriateness of regression-based modeling for quantifying interaction effects.
